# Consumer feces impact coral health in guild-specific ways

**DOI:** 10.1101/2022.10.31.514626

**Authors:** Carsten G.B. Grupstra, Lauren I. Howe-Kerr, Jesse A. van der Meulen, Alex J. Veglia, Samantha R. Coy, Adrienne M.S. Correa

**Affiliations:** BioSciences at Rice, Rice University, Houston, Texas, United States of America; The Department of Biology, Boston University, Boston, Massachusetts, United States of America

## Abstract

Microbiota from consumer feces can impact resource species in guild-specific ways. We tested the effect of fresh and heat-killed feces from corallivorous (coral-eating) and grazer/detritivorous fish on coral health and found that fresh grazer/detritivore feces, but not fresh corallivore feces, affected coral health in detrimental ways compared to heat-killed feces, suggesting that microbiota in grazer/detritivore feces were harmful. Bacterial diversity across 10 fish species suggests our experimental findings are generalizable to consumer guild: corallivore feces contained more coral-associated bacteria, and lower abundances of the coral pathogen, *Vibrio coralliilyticus*. These findings recontextualize the ecological roles of consumers on coral reefs: although herbivores support coral dominance through removal of algal competitors, they also disperse coral pathogens. Corallivore predation can wound corals, yet their feces contain potentially beneficial coral-associated bacteria, supporting the hypothesized role of corallivores in coral symbiont dispersal. Such consumer-mediated microbial dispersal as demonstrated here has broad implications for environmental management.

## Introduction

As consumers—including herbivores, predators and parasites—move through the environment and engage in consumer-resource interactions, they transmit microorganisms amongst the animals and plants with which they interact (*1*–*3*). This trophic transmission likely has important implications for microbiota assembly, as well as for resource species health and environmental acclimatization (*4*–*6*). Tropical coral reefs, for example, harbor diverse fishes that feed on benthic resource species, including stony corals and macroalgae (*7*–*9*). These fish defecate as they swim over the reef, generating a “persistent rain of feces”, containing high densities of live microbiota, that is deposited onto resident corals (*10*). This frequent deposition of fish feces–along with the microbiota they contain–is likely to affect coral health through several potential mechanisms (*11*–*15*).

Corals are filter feeders that can take up nutrients from fish waste products (*16*–*18*). Yet, fish feces also contain particulate matter derived from reef sediments and silts (*19*); when sediments land on a coral colony, it can lead to smothering and death of coral polyps, resulting in patches of mortality (lesions). Pathogenic or opportunistic bacteria in fish feces may also have negative effects on coral health (*11, 20*–*23*). For example, microbial opportunists in the feces of some grazer/detritivores can trigger lesion formation on corals (*11, 21*). This is likely because turf- and macroalgae, a major food source for these fishes, can contain diverse coral pathogens (*24*–*26*) such as members of the genus *Vibrio*; for example, *Vibrio coralliilyticus* can cause lesions or bleaching in corals (*27*–*29*).

Microbiota in fish feces may not always be detrimental to coral health, however. In fact, feces from corallivores might even support coral health through the delivery of probiotics–live, generally homologous, microorganisms that can promote the health of the recipient animal–on some coral-dominated reefs (*5, 13*). We infer this based on increased survival and growth of juvenile corals experimentally inoculated with cultured Symbiodiniaceae cells (*30, 31*), as well as increased survival of adult corals under environmental stress that were provided with mixtures of beneficial bacteria or Symbiodiniaceae cells (*32*–*38*). The feces of corallivorous fish contain high densities of potential probiotics, including live Symbiodiniaceae cells (*13, 14, 39*) and mutualistic coral bacteria including the family Endozoicomonadaceae (*40*). The deposition of feces on corals may thus affect colony health in nuanced ways depending on aspects of fecal composition including diversity of microbiota, as well as nutrition and sediment content. Studying how feces from coral reef fish in different trophic guilds—whose feces contain different microbial, nutritional and sediment compositions—affect coral health will advance coral reef ecology and inform management and conservation plans.

We tested the effect of feces from distinct consumer guilds on coral health and used heat-killed fecal controls to tease apart physical versus biological effects of contact with fecal material. We then characterized bacterial community assemblages and quantified relative abundances of the coral pathogen *V. coralliilyticus* in feces of 10 abundant consumers, ranging from obligate corallivores to grazer/detritivores, to determine whether results from the feces addition experiment are generalizable at the guild level. We tested the following hypotheses: 1) microbiota in grazer/detritivore feces, but not corallivore feces, negatively impact coral health through the development of lesions, 2) microbial communities in feces are consumer guild-specific and grazer/detritivore feces contain higher relative abundances of the coral pathogen *V. coralliilyticus*, whereas corallivore feces contain more coral-associated bacteria that are potentially beneficial.

## Methods

### Feces addition experimental design

A feces addition experiment was conducted using fragments of *Pocillopora* spp. (*41*). Briefly, fragments of 11 colonies of *Pocillopora* spp. were maintained in filter-sterilized (0.2 μm) seawater and exposed to one of five treatments for up to 22 hours (N=55 fragments from 11 colonies): fresh feces from an obligate corallivore (FC); fresh feces from a grazer/detritivore (FG); sterilized feces from an obligate corallivore (SC); sterilized feces from a grazer/detritivore (SG); and no-feces control (C; N=11 fragments per treatment; see Supplementary Methods for more detail). For the fresh feces treatments (FC, FG), we applied 100 µl fresh feces isolated from the hindgut of the obligate corallivorous butterflyfish *Chaetodon ornatissimus* (FC) or the herbivorous grazer/detritivore *Ctenochaetus striatus* (FG) directly onto each coral fragment; for the sterilized feces treatments (SC, SG), fecal pellets were first sterilized by pressure cooking in an Instant Pot (40 min using the higher pressure, high temperature settings) and then applied in the same manner (*42*). Feces for the treatments were collected from a total of ten fish representing two species, a corallivore and a grazer/detritivore (N=5 per fish species). For seven of the experimental replicates (out of 11, fresh feces and heat-killed feces treatments), feces were pooled from two corallivore or grazer/detritivore fish individuals (N=2/11 and 5/11 replicate corals for feces from two fish; see Supplementary Methods for further details). For the remaining replicates (N=4/11), feces were derived from one corallivore or grazer/detritivore individual. No treatment was applied for the no-feces control corals.

### Analysis of coral lesions and coral fragment photosynthetic efficiency

To test how microbial communities in fish feces affected coral health, we quantified the frequencies and sizes of coral lesions caused by fecal treatments, and measured the photosynthetic efficiency of each fragment. In brief, fecal pellets were removed from each fragment and photographs were taken using a dissection microscope. Dark-adapted (20 min) photosynthetic efficiency (Fv/Fm) was then measured using an imaging pulse amplitude modulation fluorometer (I-PAM; WALZ, Germany) using the default settings.

Dissection microscope photographs were analyzed in ImageJ v. 1.53f51. Each fragment was binned to one of two categories: “apparently healthy” (no change in fragment compared to before fecal application) or “lesion” (fragment contains a novel patch of bare calcium carbonate, where coral tissue died and sloughed off following fecal application). The size of each lesion was measured by determining the average polyp size on each fragment in ImageJ by counting the number of polyps in a polygon of standardized size in triplicate. Then, the size of the lesion was measured (in pixels), and the lesion size was expressed as the estimated number of polyps that had died.

Fv/Fm values were extracted from I-PAM scans of each coral fragment *post hoc*. Measurements were taken following a transect design: one measurement was taken immediately adjacent to the removed fecal pellet location, another measurement was taken halfway between the first measurement and the furthest edge away from the fecal pellet, and the final measurement was taken at the furthest edge of the fragment away from the point where feces were placed. The three measurement locations will be referred to as “Adjacent”, “Middle”, and “Far”, respectively. The effect of experimental treatment on the frequency of lesions in coral fragments (categories “healthy” or “lesion”) was tested using Chi-squared tests with the rcompanion package (v.2.4.13). Differences in lesion sizes (expressed as the estimated number of dead polyps) between treatments were tested using linear models with lm (package vegan v 1.1-28). Values were log-transformed to satisfy the assumptions of the linear model. Fragments that had complete mortality (N=4 total) were excluded from the analysis, as well as one replicate colony (all fragments/treatments) because a lesion in one of the treatments (SG treatment) was not entirely in the image and we were therefore unable to accurately assess its size. Zero values (*i*.*e*., fragments that did not have lesions) were not included in the analysis (final N: C, 0; FC, 4; FG, 8; SC, 4; SG, 4). Pairwise comparisons between all treatments were conducted using the package emmeans (v1.7.0) with a Tukey adjustment for multiple comparisons. Model assumptions were visually checked using the check_model function in the package performance v0.9.1. The effect of fecal pellets on untransformed Fv/Fm values were tested in the same way as lesion sizes, but with an interaction between the independent variables “treatment” and “distance from fecal pellet” (“Adjacent”, “Middle” or “Far”).

### Field sampling of corallivorous and grazer/detritivorous fishes, and other environmental reservoirs of coral-associated microorganisms

To test the extent to which microbial communities in fish feces—and thereby, their effects on coral health—may be broadly generalizable within fish trophic guilds, we collected additional fecal samples from each of the fish species included in the experiment (the obligate corallivore *Chaetodon ornatissimus* and the grazer/detritivore *Ctenochaetus striatus*), and other species in the same trophic guilds (grazer/detritivores, obligate corallivores), as well as facultative corallivores. Additionally, samples of corals, algae, sediments, and seawater were collected to test whether bacterial taxa in these samples were also represented in fish feces (n=5-14 per fish, coral, or algae species/genus, see Table S1). All collections were conducted in October 2020 from the back reef (1-2 m depth) and fore reef (5-10 m depth) in Moorea, between LTER sites 1 and 2 of the Moorea Coral Reef (MCR) Long Term Ecological Research (LTER) site. Obligate corallivores were defined as fish that eat corals nearly exclusively (>90% of stomach content or number of bites;, *43, 44*), and facultative corallivores were defined as fish that were observed to feed on coral for a minor to major part of their diet, while also feeding on algae or other invertebrates (∼5-∼80% of bites from corals; *12, 43*–*45*). The selected species included three additional obligate corallivore species (butterflyfishes *Chaetodon lunulatus*, “CHLU”; *Chaetodon reticulatus*, “CHRE”; and the filefish *Amanses scopas*, “AMSC”), three facultative corallivore species (butterflyfishes *Chaetodon citrinellus*, “CHCI”; and *Chaetodon pelewensis*, “CHPE”; and the parrotfish *Chlorurus spilurus*, “CHSP”), and two additional grazer/detritivore species (surgeonfishes *Ctenochaetus flavicauda*, “CTFL”; and *Zebrasoma scopas*, “ZESC”). Coral and algal samples were collected from locally abundant coral and algae genera (*46, 47*), including the corals *Acropora hyacinthus* (“ACR”), *Pocillopora* species complex (“POC”, *41, 48*), and *Porites lobata* species complex (“POR”, *49, 50*); and mixed communities of turf algae (“Turf”) as well as macroalgae in the genera *Asparagopsis* (“Asp”), *Dictyota* (“Dict”), *Lobophora* (“Lob”), *Sargassum* (“Sarg”), and *Turbinaria* sp. (“Turb”; see Table S1 for replication per species or sample category). Following collection, all samples (fish, coral, algae, sediments and water) were immediately transported on ice to the lab, where they were processed for preservation in DNA/RNA Shield (Zymo Research, CA) as described in (*13*).

### DNA Sample processing and 16S rRNA gene sequencing

DNA was extracted from all samples using the ZymoBIOMICs DNA/RNA Miniprep kit (Zymo Research, CA) according to the manufacturer’s instructions, but with a 1hr proteinase K incubation (35°C) before the lysis buffer step. Library preparation and sequencing was conducted at the Genomics Core Lab at the Institute of Arctic Biology of the University of Alaska Fairbanks. All samples were sequenced using the 16S rRNA gene V4 primers 515f (GTGYCAGCMGCCGCGGTAA) and 806rB (GGACTACNVGGGTWTCTAAT) on Illumina MiSeq using v3 2×300bp chemistry (*51*–*53*). Mock communities (HM-782D, BEI Resources, VA), extraction negatives, and plate negatives were included to facilitate the identification and removal of potential contaminants *in silico* (*54*).

Bacterial reads were processed in RStudio (version 1.1.456) through the DADA2 pipeline (version 1.11.0, *55*). The DADA2 pipeline generated a table of amplicon sequence variants (ASVs), and bacteria taxonomy was assigned using the SILVA rRNA database (version 132, *56*). Non-bacterial (*e*.*g*., mitochondrial) reads and singletons were removed. A total of 903 potential contaminant ASVs were identified and removed with the decontam package (v. 1.10.0) using the prevalence method (*54*). Libraries were rarefied to 1037 reads for the generation of Bray-Curtis distances. For visualization in stacked bar plots and calculating the number of ASVs shared between environmental pools and fish feces, all ASVs <100 reads were removed prior to rarefication to 1037 reads per sample.

To test for differences in microbial community compositions between sampling groups, PERMANOVA tests were conducted using adonis (Vegan v. 2.5-7) on Bray-Curtis distances calculated from untransformed counts tables. However, tests of multivariate homogeneity of group dispersion (using betadisper in the package Vegan v. 2.5-7) showed significant heterogeneity, which can increase the probability of Type I errors while using PERMANOVA if the sampling design is unbalanced (*57*). Because the sampling design was unbalanced—*e*.*g*., the obligate corallivore category contained four fish species with 37 samples total, while the facultative corallivore category contained three fish species with 18 samples total—tests between environmental pools (obligate corallivore feces, facultative corallivore feces, grazer/detritivore feces, coral, algae, sediments and water), and between species/sample types (*e*.*g*., feces of *A. scopas* vs. feces of *C. striatus*) were conducted separately to facilitate subsampling to create a balanced design. Thus, all the categories (environmental pool or type/species) were randomly subsampled (18 random samples per environmental pool or 5 random samples per type/species; see table S3 and S4 for sample names) prior to conducting PERMANOVA. Pairwise PERMANOVAs were conducted using the package pairwise Adonis v.0.4 using these subsampled datasets.

We tested for differences in the relative abundances of members of the family Endozoicomonadaceae, some of which are coral mutualists (*58*–*61*), between environmental pools (obligate and facultative corallivore feces, grazer/detritivore feces, corals, algae, sediment and water) using Kruskal-Wallis tests, because the residuals were not normally distributed when using linear models. Pairwise tests between environmental pools were conducted using Dunn tests in the package dunn.test v. 1.3.5. Kruskal-Wallis tests and Dunn tests were also conducted to test for differences between individual sample types/species within each environmental pool (*e*.*g*., within algae, *Turbinaria* sp. vs turf algae).

### Quantitative PCR (qPCR) of *Vibrio coralliilyticus* genes in fish feces

qPCR was conducted using vcpARTF and vcpARTR qPCR primers developed for the bacterial coral pathogen *Vibrio coralliilyticus* (*29*) and results were standardized using primers for general bacteria 967F and 1046R (*62*–*64*). Standard curves (for *V. coralliilyticus* and general bacteria) were made from a *Vibrio coralliilyticus* culture (AS008) isolated from corals collected in the Flower Garden Banks National Marine Sanctuary (northwest Gulf of Mexico). Sequencing of the full-length 16S gene region of bacterial DNA with primers 8F and 1513R resulted in 97.5% percent identity with *V. coralliilyticus* strain U2 (accession MK999891.1) with 100% query cover. The primers vcpARTR and vcpARTF were used to amplify metalloprotease genes and the amplicon was then cleaned using a genejet PCR purification kit (Thermo Fisher, MA); resultant DNA concentrations were acquired using Qubit (Thermo Fisher, MA). A standard curve was made using serial dilutions from 10^9^ to 10^0^ gene copies per µl template; the standard curve for *V. coralliiyticus* primers vcpARTR and vcpARTF had an efficiency of 106.2%; the standard curve for general bacteria primers 967F and 1046R had an efficiency of 92.3%. Sanger sequencing of gene fragments amplified using vcpARTF and vcpARTR primers from DNA extracted from *C. striatus* feces resulted in a top hit against a *V. coralliilyticus* strain P4 metalloprotease gene (accession JQ345042.1) with an e-value of 3×10^−12^ (query cover 58%, percent identity 85%). Amplification of the *V. coralliilyticus* target gene at <30 cycles was counted as a positive detection, which roughly corresponded to amplification of the 10^2^ gene copies µl^-1^ standard (Ct=30.4). Delta cycle threshold (dCT) values were calculated by subtracting the cycle threshold at which the signal from general bacteria primers was detected from the cycle threshold at which *V. coralliilyticus* was detected. Higher dCT values indicate lower relative abundances of *V. coralliilyticus*. Differences in dCT values between fecal samples of each trophic guilds were tested with an ANOVA, followed by Tukey tests for multiple comparisons (using a Tukey correction). The assumptions for the test were visually assessed.

### Data availability

Sequence reads have been uploaded to NCBI’s seqence read archive (SRA) under accession [XXXXX]; all other data have been included as supplementary data files. R code to replicate the analyses has been deposited on GitHub: https://github.com/CorreaLab/graco.

## Results

### Microbiota in grazer/detritivore feces increased coral lesion frequencies and sizes

Fecal treatments resulted in the formation of lesions or complete mortality in some coral fragments, while other replicates remained visually healthy (Figure 1A, S1; X^2^=20.93, p-value=0.0005; Supplementary Data File 1). Fresh grazer/detritivore feces resulted in the formation of lesions or complete mortality in all replicates (N=9; 2, respectively), while the fresh corallivore feces treatment resulted in lesions or total mortality in only some fragments (N=4; 1, respectively) and most fragments remained visually healthy (N=6). Effects of both sterile feces treatments on fragment health were comparable to those caused by fresh corallivore feces (i.e., lesions or total mortality in only some fragments; sterile corallivore – N=4 and 0; sterile grazer – 5 and 1). All control (no feces treatment) fragments remained visually healthy during the experiment.

**Figure 1.**
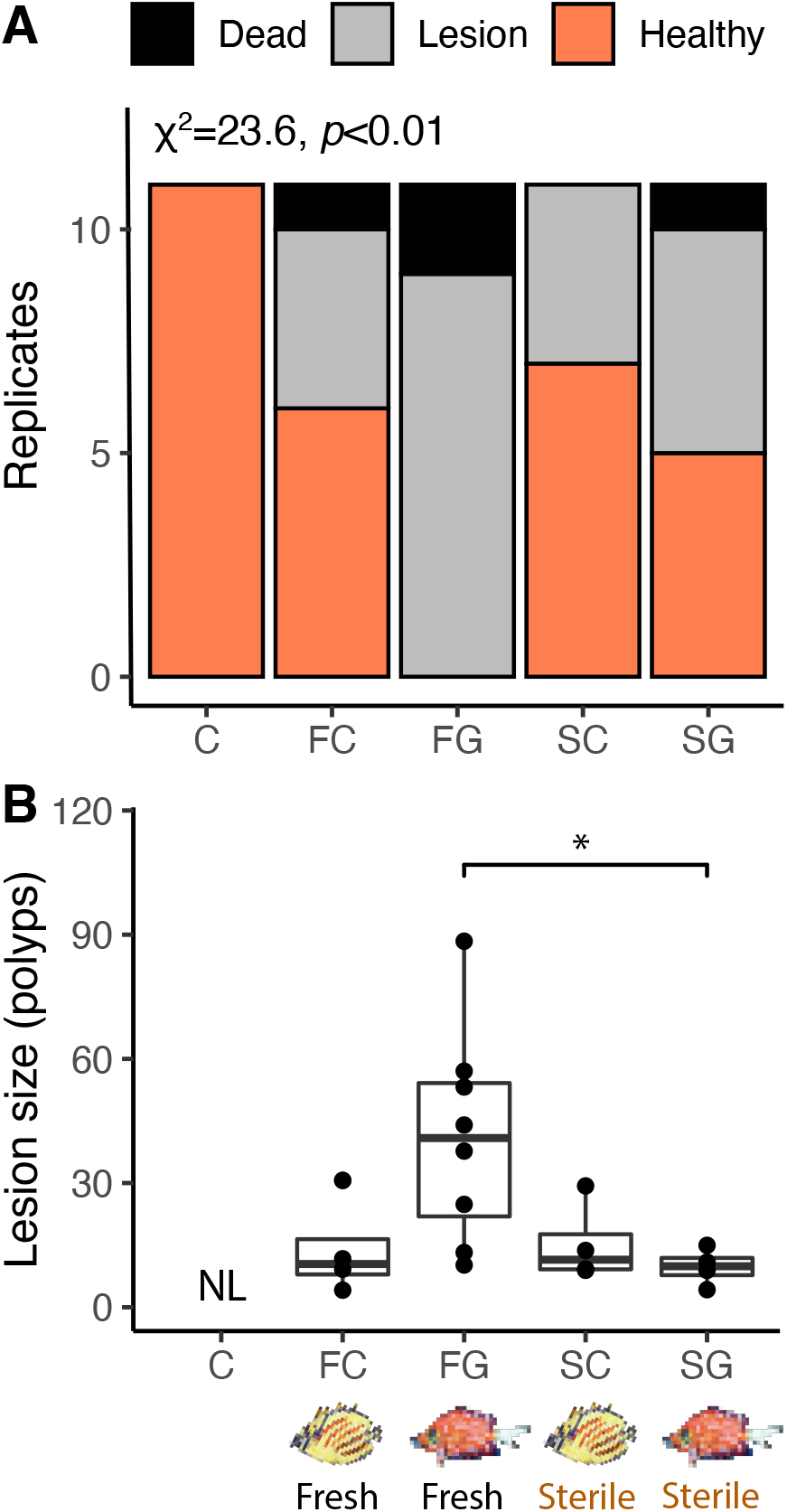
Fish feces donor species (corallivore or grazer/detritivore) influenced coral tissue integrity (A) and lesion size (B). (A) Fresh grazer feces caused lesions or mortality in all replicates (N=11/11), whereas fresh corallivore feces only caused lesions or mortality in 46% of replicates (N=5/11). Sterilized feces caused lesions in only 45% (N=5/11; sterile corallivore feces) to 55% (N=6/11; sterile grazer feces) of replicates (overall chi-squared test results: X^2^=20.93, p-value=0.0005). (B) Fresh grazer feces application caused lesions that were, on average, 4.2 times larger than sterile grazer feces (LM results: Treatment F=4.38, Df=3, p=0.02; pairwise comparisons FG-SG: estimate=1.322; adjusted p=0.031, see Table S2 for individual comparisons). Together, these findings show that grazer feces can be biologically harmful to corals upon contact, whereas corallivore feces are not. “NL”: No lesions observed.

Lesion sizes caused by fecal pellets also differed significantly among treatments (Figure 1B; Df= 3, F=4.38, p=0.02; Supplementary Data File 2). Pairwise comparisons revealed that mean (± SD) lesion sizes in the fresh grazer feces treatment (41.1 ± 25.8 polyps) were, for example, 4.2 times larger than those in the sterile grazer feces treatments (9.76 ± 4.46 polyps; est= 1.32; p=0.031, see Table S2 for all pairwise comparisons). Lesions caused by grazer feces were also three times larger than those caused by fresh corallivore feces (13.9 ± 11.6 polyps), but this pairwise comparison was not significant, potentially due to the low number of lesions caused by fresh corallivore feces (N=4). Lesion sizes caused by fresh corallivore feces did not differ from sterile corallivore feces (15.3 ± 9.6 polyps). Finally, there was no effect of fecal treatment on coral photosynthetic efficiency (Figure S2; LM results: treatment*distance: Df=8, F=0.045, p=0.999; treatment: Df=4, F=2.27, p=0.065; distance: Df=2; F=0.53, p=0.588).

### Bacterial communities differed between environmental pools

To determine whether the microbiota in feces of the corallivore and the grazer/detritivore fish species used in the feces addition experiment are representative of microbiota at the guild-level (and may thus have similar effects on coral health), we quantified the bacterial diversity in feces of three fish trophic guilds (four species of obligate corallivore, three species of facultative corallivore, and three species of grazer/detritivore; see Table S1 for replication; Supplementary Data File 3-5). Bacterial communities in fish feces were also compared to those in other corals, algae, sediment and water (see Table S1 for replication) to characterize the extent to which fish feces contain the microbial taxa associated with their main food sources (all samples combined, N=161).

Quality filtering and removal of singletons and potential decontaminant ASVs resulted in 161 sequencing libraries (excluding negatives and mock communities) with a mean read depth of 14,054 reads per sample. A total of 16 samples were discarded because the libraries had <1,000 reads after processing. These included the following samples: 5 *Acropora hyacinthus, 5 Porites lobata* spp., 4 *Pocillopora* spp., 1 sediment and 1 water sample). Bacterial communities in all environmental pools (*i*.*e*., obligate and facultative corallivore feces, grazer/detritivore feces, corals, algae, sediment and water) were significantly different from one another (Figure 2; overall PERMANOVA results: R^2^=0.154, F=3.725, p<0.001, pairwise PERMANOVA results: R^2^>0.064, F>2.3, Bonferroni-adjusted p-values<0.03; see Table S3). Bacterial communities associated with many of the individual species/sample types also differed from each other (*e*.*g*., bacteria in feces from *Chaetodon ornatissimus* versus those in *Chlorurus spilurus*; Figure 2; overall test results R^2^=0.409, F=2.903, p<0.001; see Table S4). Out of all pairwise comparisons among species/sample types (210 pairwise comparisons total), most (188, 89.5%) were significant, including all pairwise comparisons between obligate corallivore species and grazer/detritivore species. Out of the pairwise comparisons that did not differ significantly (22, 10.5%), most comparisons (13) were between different types of algae, sediment, and the coral *Acropora hyacinthus*. Interestingly, two obligate corallivores (CHOR, CHRE) and one facultative corallivore (CHPE) did not differ significantly from *Porites lobata* species complex microbiomes, suggesting overlap in microbial community compositions (but *Pocillopora* species complex samples did differ significantly from these fish feces samples). Additionally, several of the obligate and facultative corallivores did not differ significantly from each other (CHOR and CHPE, CHLU and CHOR, CHPE and CHRE, CHOR and CHRE; Table S4). Comparisons between two grazers and between a grazer and a facultative corallivore were also insignificant (CHSP and CTST, CTST and ZESC; Table S4).

**Figure 2.**
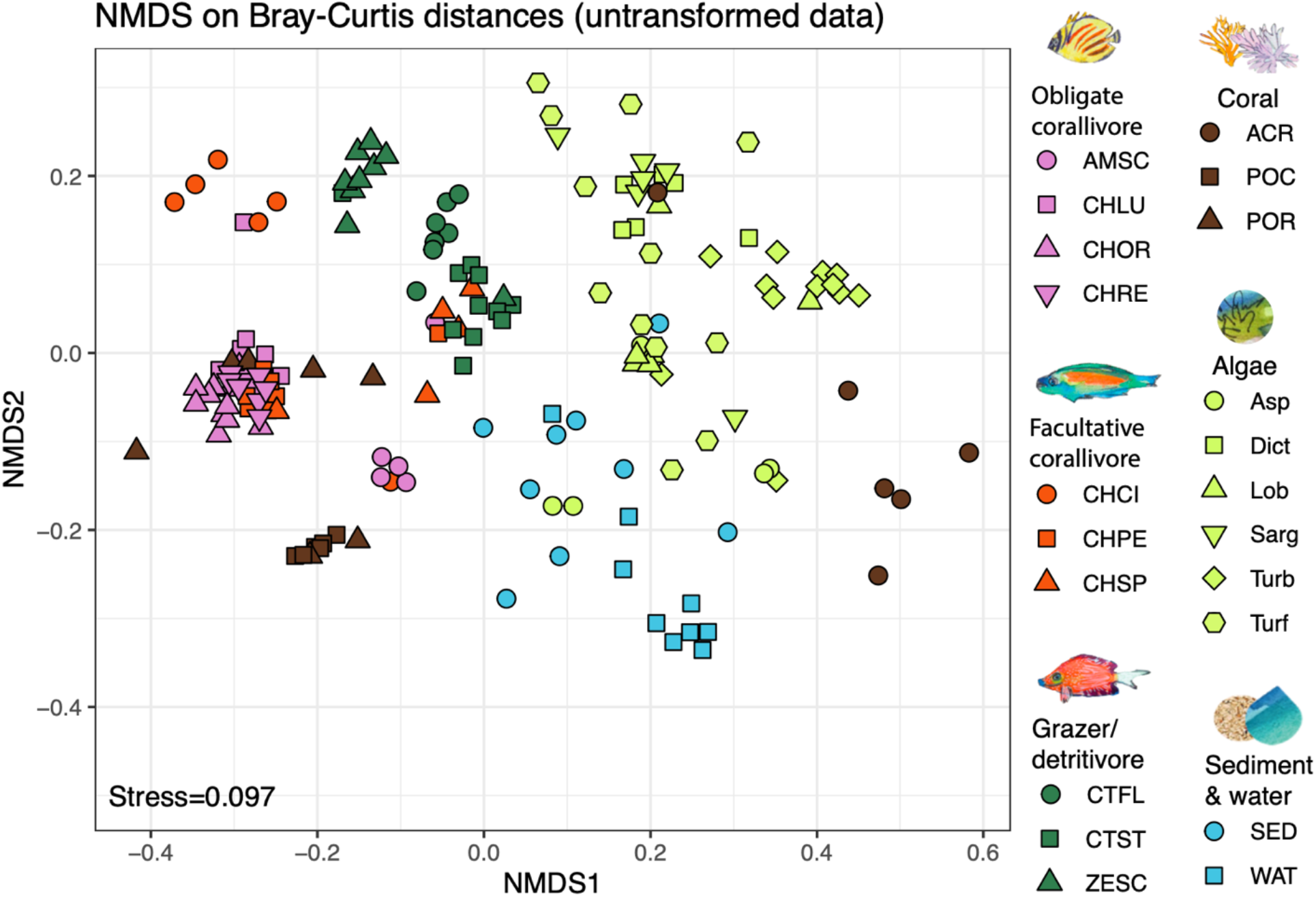
NMDS of bacterial communities based on the 16S rRNA gene in coral reef-associated environmental pools: obligate and facultative corallivore feces, grazer/detritivore feces, corals, algae, sediment and water. Bacterial communities in all environmental pools differed significantly from each other (overall PERMANOVA results: R^2^=0.154, F=3.725, p<0.001, pairwise PERMANOVA results: R^2^>0.064, F>2.3, Bonferroni-adjusted p-values<0.03; see Table S3). Bacterial assemblages also significantly differed between most species/sample types (overall PERMANOVA results: R^2^=0.409, F=2.903, p<0.001; see Table S4 for individual comparisons). PERMANOVA results based on Bray-Curtis distances calculated from untransformed sequencing libraries with taxa <100 reads removed and then rarefied to 1,037 reads per sample. See methods for species abbreviations.

### Consumers disperse bacterial taxa associated with the resource that they consume

Feces of individual obligate corallivore species contained up to 88.6% of the distinct ASVs found in *Porites lobata* species complex (*C. reticulatus*; 225 of 254 ASVs), and up to 75.6% of ASVs found in *Pocillopora* species complex (*A. scopas*; 90 of 119 ASVs; Figure 3, Table S6). By comparison, grazer/detritivore feces contained only up to 9.1% of distinct ASVs associated with *Porites lobata* species complex (*C. striatus*; 23 of 254) and up to 5.0% of the ASVs identified in *Pocillopora* species complex (*Z. scopas*; 6 of 119). ASVs associated with *A. hyacinthus* were much less frequently identified in corallivore feces (up to 3.1%; *A. scopas*), but these corals were relatively uncommon at our study site during the sampling period, and are not frequently consumed by corallivores (*13*).

**Figure 3.**
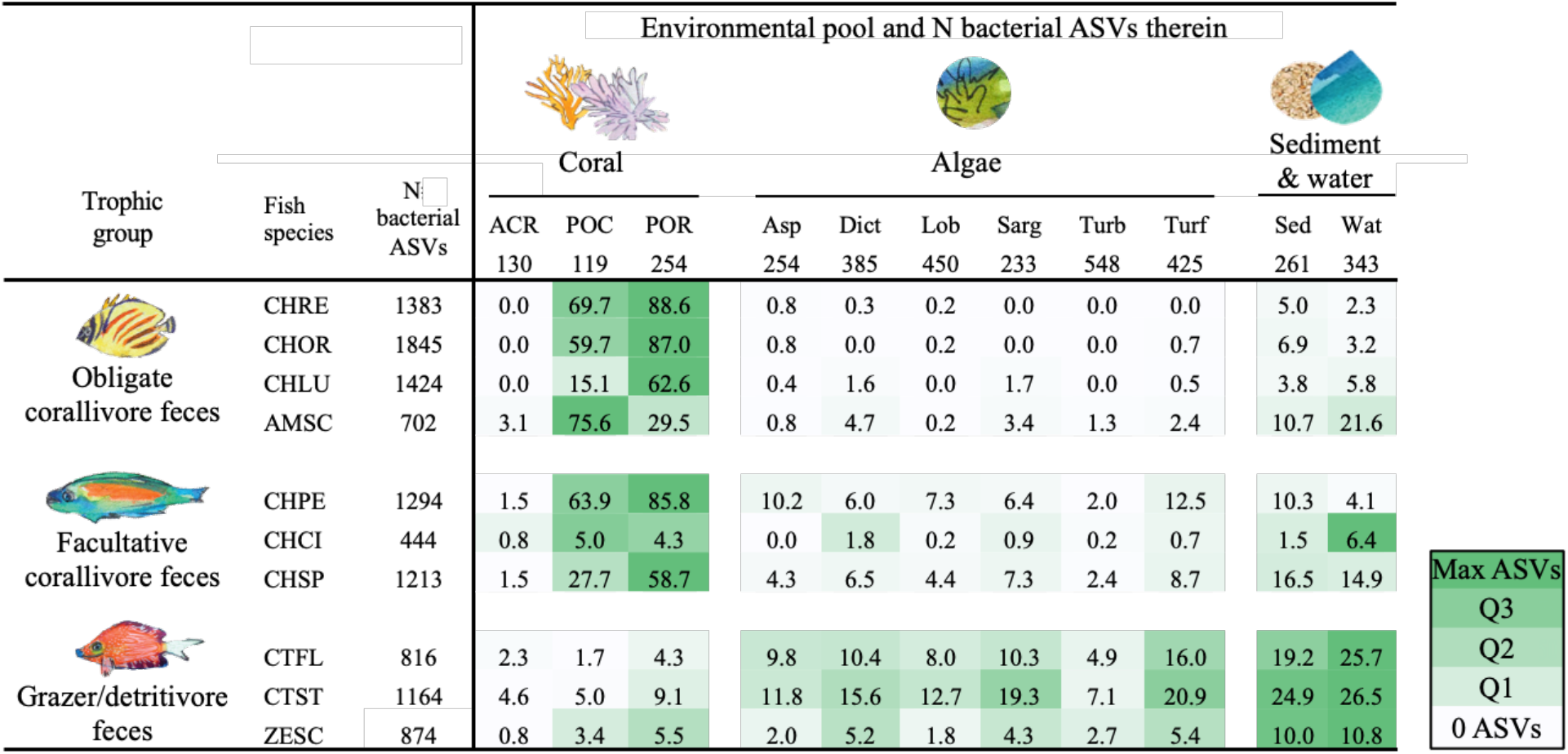
Coral reef fish disperse bacterial taxa associated with the resource that they consume. Listed are the fractions (%) of bacterial ASVs associated with corals, algae, and sediment and water that were also identified in the feces of each fish species. Darker shading indicates higher representation (%) of bacterial taxa; shading was scaled for each row (*i*.*e*., each consumer species) separately. All taxa with less than 100 reads were removed, and all libraries were rarified to 1,037 reads. All samples per species or environmental pool were pooled. “N bacterial ASVs” indicates the total number of unique ASVs in each environmental pool or fish species after rarefaction. See Table S5 for the number of shared ASVs. See methods text for species abbreviations.

Grazer/detritivore feces contained ASVs that were associated with different types of algae (Figure 3, Table S6). For example, 20.9% of ASVs associated with sampled turf algae were identified in feces of *C. striatus* (69 of 425 ASVs), whereas only up to 2.4% of turf algae ASVs were found in obligate corallivore feces (*A. scopas*; 10 of 425 ASVs). In addition, grazer/detritivore feces contained higher numbers of the ASVs associated with all other categories of algae than did feces of obligate or facultative corallivores (Figure 3, Table S6). Grazer/detritivore feces also contained more of the ASVs found in sediment and water samples (*C. striatus*; 24.9 %, 65 of 261 ASVs and 26.5%, 91 of 343 ASVs), than did other trophic guilds. Facultative corallivore feces contained a mix of ASVs associated with corals and algae.

For example, the facultative corallivore *Chaetodon pelewensis* contained 85.8% of the ASVs found in *Porites lobata* species complex, as well as up to 12.5% of the ASVs associated with algal samples. Interestingly, feces of the facultative corallivore *C. citrinellus* contained fewer of the ASVs associated with corals or algae than other members of its guild; fecal samples of this species contained only 0.8-5% of ASVs associated with coral samples and 0-1.8% of ASVs associated with algal samples.

### Relative abundances of the coral mutualist Endozoicomonadaceae were higher in feces of obligate corallivores than grazer/detritivores

Relative abundances of Endozoicomonadaceae reads in amplicon sequencing libraries significantly differed among environmental pools (Figure 4; Kruskal-Wallis test, X^2^=94.5, df=5, p<0.001). Pairwise tests demonstrated that mean (±SE) relative abundances of Endozoicomonadaceae reads were significantly higher (|Z|>2.2, adjusted p-values<0.04, see Table S5) in corals (43.3±32.8%), obligate corallivore feces (23.9±9.5%), and facultative corallivore feces (25.4±10.5%) than in grazer feces (12.8±10.5%), algae (3.3±2.7%), or sediment and water (5.3±5.2%). Relative abundances of Endozoicomonadaceae reads were also higher in feces of the obligate corallivore *C. ornatissimus* than the grazer/detritivore *C. striatus*, which were the two species used for the feces addition experiment (Kruskal-Wallis test tesults: X^2^=13.295, df=1, p=0.0003). Relative read abundances of Endozoicomonadaceae further differed between species/sample types within environmental pools or trophic guilds (Table S6). For example, relative abundances of Endozoicomonadaceae reads in feces differed among obligate corallivore species; they were lower in feces of *A. scopas* than in those of *C. ornatissimus* and *C. reticulatus* (Kruskal-Wallis test results: X^2^=8.892, df=3, p=0.031; Dunn test results |Z|>2.5, p<0.05) see Table S6).

**Figure 4.**
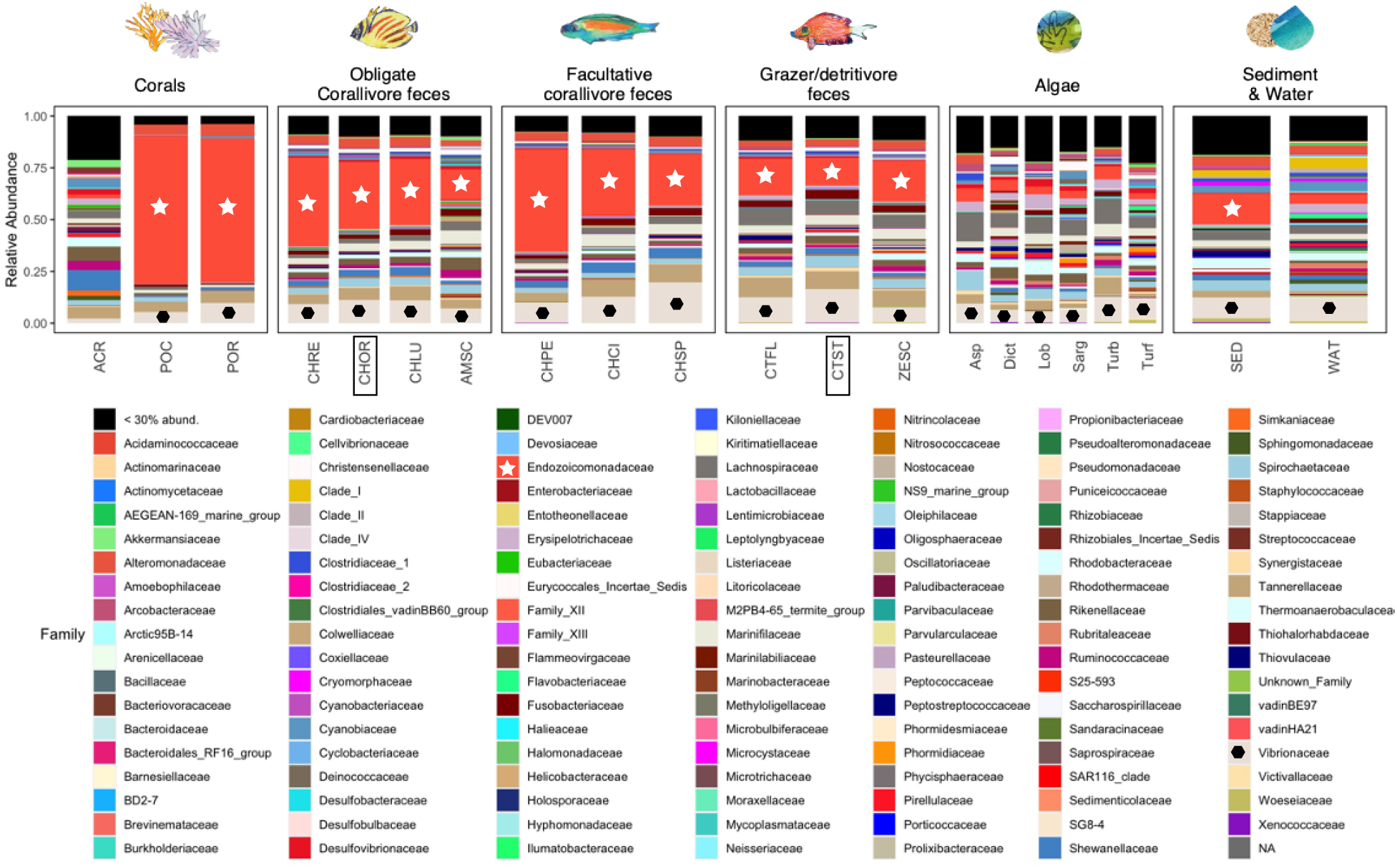
Stacked bar plots illustrate that relative abundances of bacterial families differ among environmental pools, consumer trophic guilds and species. Stacked bar plots are based on sequencing libraries subsampled to 1,037 reads. Notable families include Endozoicomonadaceae (crimson, marked with white star where possible), which are coral mutualists and (in this study) are abundant in the feces of obligate and facultative corallivores (see Table S6 and S7 for pairwise comparisons of relative abundances). The family Vibrionaceae (sand colored, marked with black polygon where possible), which includes some notable coral pathogens, are present in all sample types. Fish species used in the feces addition experiment (Figure 1) are indicated with boxes. All taxa with less than 100 reads were removed, all libraries were rarified to 1,037 reads, and bacterial families that represent <30% of total reads were collapsed (“<30% abund.”). “NA”: No family assignment.

### *Vibrio coralliilyticus* relative abundances were higher in feces of grazer/detritivores than in corallivore feces

The *Vibrio coralliilyticus* metalloprotease gene was amplified using qPCR of DNA extracted from feces of each sampled fish species (Figure 5). Using this method, *V. coralliiyticus* genes were identified (Ct<30; see Methods) in 54% (N=20/37) of obligate corallivore feces, 50% of facultative corallivore feces (N=9/18), and in 65% (N=20/31) of grazer/detritivore feces. Relative abundances of *V. coralliilyticus* genes differed between fish guilds (ANOVA: Df=2, F=16.14, p<0.001; Supplementary Data File 6). A Tukey post-hoc test revealed that relative abundances of *V. coralliilyticus* genes were significantly higher in grazer/detritivore feces than in obligate corallivore feces (Figure 5; difference = 4.61, p<0.001) and facultative corallivore feces (Figure 5; difference = 2.44, p=0.046).

**Figure 5.**
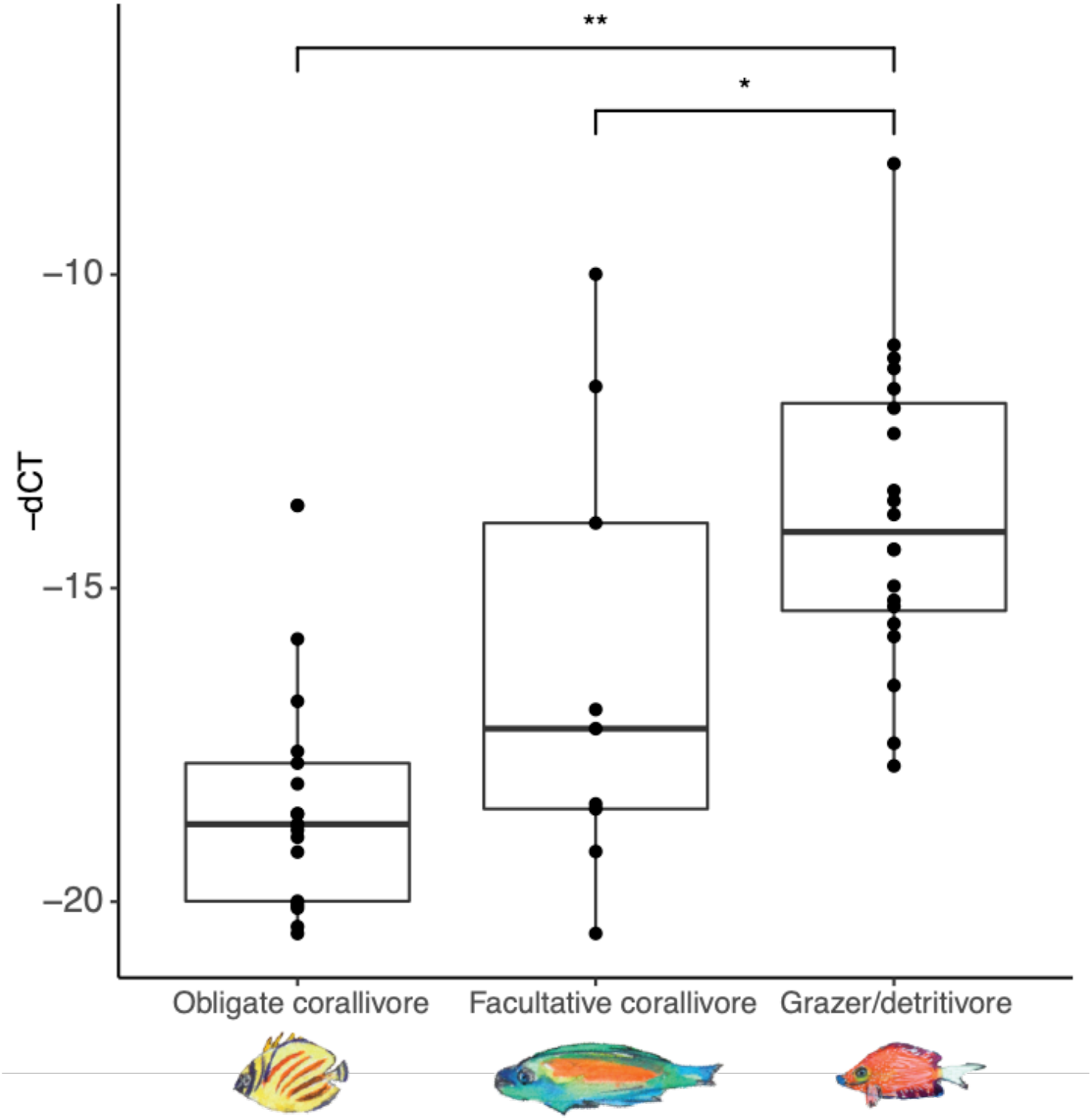
Relative abundances of *Vibrio coralliilyticus metalloprotease* genes were higher in grazer/detritivore feces than obligate corallivore and facultative corallivore feces (overall ANOVA results: F= 16.14, p<0.001). Delta cycle threshold (dCT) was calculated by subtracting the cycle threshold for the general bacteria assay from the *V. coralliilyticus* assay; therefore, samples with lower dCT values contained higher relative abundances of *V. coralliilyticus*. Note that the y-axis has been inverted to make interpretation of the results more intuitive. P-values: * <0.05; ** <0.001.

## Discussion

The production of waste products is ubiquitous across the animal kingdom; this waste is a key component of nutrient cycles as well as an important way in which microbiota are transported throughout environments (*5*). Animal diets determine the composition of waste products, including their associated microbiota, and thus how the waste may affect other organisms in the environment (*1, 40, 65*). Testing how feces derived from animal species in diverse trophic guilds affect cohabitating organisms can inform agriculture, ecosystem management, and restoration. Here, we demonstrate that feces from distinct trophic guilds of coral reef fish—grazer/detritivores and corallivores—affect the health of reef-building corals in distinct ways. Microbiota in grazer/detritivore, but not corallivore, feces affect coral health in detrimental ways. Subsequent characterization of bacterial diversity in grazer/detritivore and corallivore feces demonstrated that these communities were trophic guild-specific, suggesting that findings from the fecal addition experiment presented here are generalizable to trophic guild. Together, these findings highlight an underexplored, indirect result of consumers on resource species, which has potentially important ramifications for ecosystem health and functioning.

### Microbiota in feces of grazer/detritivore fish, but not corallivorous fish, are detrimental to coral health

Intact coral reef ecosystems are characterized by high abundances of diverse fishes that release feces, containing up to 2.6×10^11^ bacterial cells g^-1^ dry weight of feces, as they move through their territories (*7*–*10*). This frequent deposition of feces is likely to affect coral microbiota assembly, and thereby, colony health. The feces addition experiment presented here demonstrated that fresh feces from the grazer/detritivore *C. striatus* caused detrimental effects to coral health (higher frequency of lesions and larger lesions) compared to sterilized feces from the same species (Figure 1B, 2; Table S2), suggesting that microbial activity in fresh grazer/detritivore feces is driving these detrimental effects to coral health. This may be caused by higher relative abundances of pathogens like *V. coralliiyticus* in *C. striatus* feces, or the presence of other potential pathogens that are associated with the algae that they eat (Figure 3, Table S6;, *24*–*26*). By comparison, fresh feces from the obligate corallivore *C. ornatissimus* did not cause significantly larger lesions on corals than sterilized feces from the same species, suggesting that the microbiota in feces of this species were not detrimental to coral health (Figure 1, 2).

To test whether the findings from the feces addition experiment are broadly generalizable, we compared the bacterial assemblages in feces from 10 fish species in three trophic guilds: obligate corallivores, facultative corallivores, and grazer/detritivores. Bacterial communities significantly differed between trophic guilds, as well as between individual species within trophic guilds (Figure 2, 4, Table S3, S4). While grazer/detritivore feces contained more bacterial taxa associated with algae (Figure 3, S5), most corallivore feces contained a higher percentage of coral-associated bacterial ASVs (Figure 3, S6) and higher relative abundances of the coral mutualist family Endozoicomonadaceae (Figure 4, Table S6, S7). Notably, grazer/detritivore feces also contained higher abundances of *V. coralliiyticus* (Figure 5), which is a known coral pathogen (*27*). Together, these lines of evidence strongly suggest that bacterial assemblages in fish feces are an important determinant of how these feces affect coral health, and feces from different fish trophic guilds are likely to impact coral health in distinct ways.

Herbivorous fishes, including grazer/detritivores, are recognized as important contributors to coral reef resilience (*9, 66, 67*). By removing algal biomass, herbivores exert top-down control on algal communities, modulating the competition for space between corals and algae, which in turn promotes coral recruitment and growth (*67*–*69*). Herbivore activity thereby increases coral reef resilience under anthropogenic pressures and reduces the chance of phase shifts from coral- to algae-dominated states (*70*). Our research, as well as two previous studies (*11, 21*), add an additional dimension to this relationship: herbivores can also disperse pathogens in their feces that cause lesions in coral tissues. More work is needed to test the extent to which different (herbivorous) fish species contribute to the dispersal of viable coral pathogens (*5*).

### Fecal pellet integrity and potential macrosymbiont interactions

Corals may acquire nutrients (*16*–*18*) and beneficial or harmful microbiota from fish feces (*11, 13*–*15, 21, 39, 71*). As shown here (Figure 1), particulate matter in fish feces may also sometimes smother coral polyps, resulting in (partial) mortality (*19, 72*). One factor that may influence how fish feces affect coral health includes the consistency and integrity of the fecal pellet. In this fecal addition experiment, feces placed on coral fragments mimicked intact feces, yet *in situ*, fecal pellets sometimes fall apart in the water column (*13*), resulting in the release of silt to rice grain-sized particles. Such particle sizes may be less likely to cause smothering of coral polyps, and might be more readily removed or even taken up and digested by corals (*73, 74*); this should be experimentally tested. We did not examine whether coral fragments in this experiment consumed parts of fecal pellets; further, a lack of data on coprophagy behavior by *Pocillopora* spp., and corals in general, limits estimates of the likelihood and frequency with which corals may consume fish feces and associated microbiota.

We also observed *Trapezia* crabs, which are invertebrate macrosymbionts of branching corals, breaking up and consuming corallivore feces applied to coral fragments *in situ* as well as in the lab (but note that *Trapezia* crabs were removed from coral fragments in this experiment). Through such activity, macrosymbionts may in some cases prevent lesions from forming when feces fall directly on coral surfaces. Empirical tests of the extent to which fecal particle size and macrosymbiont activity influence the outcomes of fish fecal contact with corals are needed.

### Potential for beneficial effects of coralivorous fish feces on coral health

Consumers in diverse ecosystems fulfill important roles regarding the dispersal of microbiota (*3, 5, 75, 76*). Through the dispersal of microbiota, consumer activities affect the assembly of microbiota in resource species (*5*). Here, we show that feces from reef fish contain a high diversity of bacterial taxa (ASVs) that are associated with their food sources. For instance, obligate corallivore feces contained up to 88.6% of the bacterial ASVs found in a locally abundant coral species (*Porites lobata* species complex; Figure 3). We therefore posit that the presence and activity of corallivores promotes the dispersal of bacterial taxa associated with corals. Based on experimental demonstration (this study) of the impact of microbial activity on coral health, we infer that at least some bacteria in fish feces are viable. Empirical tests of this (e.g., through culturing, or membrane permeability-based stains and cell sorting or viability PCR;, *5*) will help reveal the extent to which passage through fish digestive tracts affects the viability of microbiota dispersed in the feces of reef fish.

While some corals provide their offspring with key bacteria (*77*–*79*), most corals acquire at least some, or most, microbial taxa *de novo* each generation from the environment (*80*–*82*). In many coral species, juveniles may take up diverse microbial taxa from environmental pools, followed by a winnowing process in which microbial assemblages are “refined” (*81, 83*–*87*). Corallivore feces may represent hotspots of homologous bacteria and facilitate the acquisition of locally beneficial microbiota by corals and other hosts. While the transfer of beneficial bacteria between coral colonies via fish feces has not been demonstrated, earlier studies have shown that anemones in the genus *Aiptasia*—which are closely related to corals—can take up Symbiodiniaceae cells from the feces of obligate corallivorous butterflyfishes (*14*); coral juveniles have also been demonstrated to initiate symbiotic partnerships with Symbiodiniaceae cells from the feces of giant clams (*15*). Hence, it is likely that juvenile corals can acquire beneficial bacteria from fish feces as well.

The dispersal of coral-associated bacteria by corallivores may also be beneficial to stressed adult coral colonies. Environmental stress can result in disruption of the symbiosis between coral animals and their populations of microorganisms (*88, 89*, but see *90, 91*), yet experimental inoculations with engineered “probiotic” cocktails consisting of cultured coral-derived bacteria and Symbiodiniaceae have increased photosynthetic efficiency and colony health in thermally stressed corals within hours to days after inoculation (*33*–*37*). Such probiotics may promote coral health by providing nutrition, mitigating toxins, deterring pathogens, and stimulating development (*35*). For example, inoculations with microbiota from heat tolerant coral colonies were shown to reduce bleaching rates in recipient heat-susceptible colonies of the coral species *Pocillopora* sp., and *Porites* sp. in Thailand (*32*). Given that corallivore feces contain many of bacterial ASVs associated with locally abundant corals, and previous work has shown that such feces contain high densities of live coral-associated Symbiodiniaceae cells (*13, 14, 39*), corallivore feces may in some cases be expected to have a similar “stabilizing” effect on the partnership between corals and their microbiomes. This is especially likely given reports that some corallivorous fish preferentially feed on heat tolerant corals during bleaching events (*92, 93*), thereby potentially transferring microbiota associated with stress-resistant colonies to surrounding stress-susceptible colonies. Empirical studies are needed to explicitly test this hypothesis. Additionally, future studies may examine how repeated applications of feces affect coral health over longer time scales and under environmental stress, and how exposure to feces affects other measures of coral health, including tissue content (e.g. lipid, carbohydrate and protein stores, chlorophyll content) and gene expression.

## Conclusions

Our results indicate that feces of grazer/detritivores, but importantly, not corallivores, have negative effects on coral health when they come in direct contact with colonies. Feces from grazer/detritivore fish contain higher proportions of bacterial ASVs associated with locally abundant algae, as well as higher relative abundances of some coral pathogens, than feces from corallivores. These findings add a new dimension to our understanding of how fish trophic guilds influence coral health: although grazer/detritivores support coral dominance through removal of algal competitors, they also disperse pathogens that can harm coral health. Corallivore predation can wound corals, yet they disperse bacteria in their feces that may be beneficial to corals, supporting the hypothesized role of corallivores in coral-associated microbiota dispersal. Future studies may test how consumers in additional fish trophic guilds affect coral health, as well as disentangle how future climate scenarios will modulate interactions between corals and fish feces. More broadly, studying how different trophic guilds of consumers contribute to the dispersal of microbiota can aid ecosystem management and conservation, for example through the identification of trophic guilds that may promote or harm resource species in different environmental contexts.

## Acknowledgements

We thank Dr. Kory Evans, JoJo West, Sean Trainor, Moana Le Rohellec and Dennis Conetta for their help with fish collections and surveys. We thank Dr. Amanda Shore for providing the *Vibrio coralliilyticus* cultures used for the qPCR standard curve, and Dr. Deron Burkepile for providing some materials used in this work. We thank Julianna Renzi for discussion regarding coral ectosymbionts that informed the interpretation of some of our findings. Lastly, we would like to thank Cyrus Washington, Jake Sperry, and Sara Emami for their contributions to lab work, and Daniel Gorczynski for feedback on the manuscript. This research was completed under permits issued by the Territorial Government of French Polynesia (Délégation à la Recherche) and the Haut-Commissariat de la République en Polynésie Francaise (DTRT) (Protocole d’Accueil 2013–2019), and we thank the Délégation à la Recherche and DTRT for their continued support. All samples were collected under approval of the institutional animal care and use committee (IACUC) at Rice University [IACUC-21-019]. This work represents a contribution of the Moorea Coral Reef (MCR) LTER Site (NSF OCE 16– 37396). This study was made possible through U.S. National Science Foundation awards (OCE #2145472 and #1635798) and Rice University start-up funds to A.M.S.C. and fellowships awarded by the Wagoner Foreign Study Scholarship Program to C.G., J.M., L.H.K., and A.V. The Kirk W. Dotson Endowed Graduate Fellowship in Ecology and Evolutionary Biology also supported C.G., L.H.K. and A.V. in completing this work.

