## Supplementary Materials for "Consumer feces impact coral health in guild-specific ways"

### Supplementary methods

The fecal addition experiment was conducted three times (14 replicates total): an initial iteration was performed in October 2020 (four replicates) and two additional iterations in June 2021 (5 replicates each). For the initial iteration, four colonies of the cryptic coral genus *Pocillopora* spp. were collected from the fore reef at ~10 m depth; these colonies were later identified to be members of the species *Pocillopora meandrina* following methods in (*41*). After 24 h, all corals were cut into 5 fragments and left to acclimate for 48 hours, and any macrosymbionts (e.g. *Trapezia* crabs, shrimp) were removed. All fragments were transferred to glass jars containing ~500 ml of 100 KDa-filtered (sterile) seawater, photographed under a dissection microscope with 3.5-180X zoom (AmScope SM-1TSZZ-144S-10M), and the fluorescence and absorption spectra measured using an imaging-PAM (Waltz, Germany) following a 20-minute dark adaptation.

We then blindly assigned each jar containing a coral fragment to one of five treatments, so that one fragment from each coral colony was assigned to each treatment (4 replicate colonies x 5 treatments = 20 coral fragments in separate jars): fresh feces from a corallivorous butterflyfish (FC); fresh feces from a grazer/detritivore (FG); sterilized feces from a corallivorous butterflyfish (SC); sterilized feces from a grazer/detritivore (SC); no-feces control (C). For the fresh feces treatments (FC, FG), we applied 100 µl fresh feces isolated from the hindgut of the butterflyfish *C. ornatissimus* (FC) or the grazer *C. striatus* (FG) directly onto each coral fragment; for the sterilized feces treatments (SC, SG), fecal pellets were first autoclaved for 40 minutes at 120 ºC and then applied in the same manner. The feces used in the experiment (fresh and sterile) were subsampled for DNA extractions by preservation in DNA/RNA shield (Zymo Research, CA). No manipulation was conducted for the no-feces control corals. After ~22 hours, all fragments were photographed under the dissection microscope before and after the removal of the fecal pellet, and the fluorescence and absorption spectra were measured using imaging-PAM following a 20-minute dark adaptation.

The same experimental design was used for the second and third iteration, but with minor modifications: first, all coral colonies used in the experiment were sampled before collection and their species identity was determined as *Pocillopora verrucosa* using restriction fragment length polymorphism (RFLP) analysis, as outlined in (*1*). For these iterations, five replicate colonies were used instead of four. Additionally, fecal treatments in these iterations were composed of feces from two individuals per fish species that were mixed prior to application (instead of using feces from a single fish individual per treatment).

The third iteration of the experiment was ended prematurely (after ~15 hours instead of ~22 hours) because fragments (including control fragments) from three of five colonies were exhibiting tissue loss. All data for the entire set of samples (all treatments) for these three colonies were discarded and not included in subsequent analyses. For all subsequent analyses, replicates from all iterations were analyzed together.

*Bacterial culturing*

To ensure that the sterilization method for the sterile feces treatments worked as intended, we streaked agar plates with subsampled material of each fecal treatment in iteration 2 and 3 of the experiment. Briefly, sterile plates (Difco™ Marine Agar 2216, 5.51%) were briefly swabbed with a rice grain-sized mixture of fresh or sterile feces from of *Chaetodon ornatissimus* or *Ctenochaetus striatus*. Culture plates were monitored for bacterial growth for five days and transferred to fresh agar plates. Bacterial growth was observed in 10 of 10 agar plates with fresh fecal samples and one of eight agar plates with sterilized fecal samples.

### Supplementary Tables

**Table S1** Summary of samples collected and successfully sequenced (>1,000 reads) in this study for the characterization of bacterial community composition in environmental reservoirs (fish feces, corals, algae, sediment and water).

| Sample category and type | Code | Back Reef | | Fore Reef | | **Total** | |
|---|---|---|---|---|---|---|---|
| | | Collected | >1,000 reads | Collected | >1,000 reads | **Collected** | **>1,000 reads** |
| **Obligate corallivores** | | | | | | | |
| *Amanses scopas* | AMSC | - | - | 5 | 5 | **5** | **5** |
| *Chaetodon lunulatus* | CHLU | 6 | 6 | - | - | **6** | **6** |
| *Chaetodon ornatissimus* | CHOR | 6 | 6 | 8 | 8 | **14** | **14** |
| *Chaetodon reticulatus* | CHRE | - | - | 8 | 7 | **8** | **7** |
| Category sum | | 12 | 12 | 21 | 20 | **33** | **32** |
| **Facultative corallivores** | | | | | | | |
| *Chaetodon citrinellus* | CHCI | 6 | 6 | - | - | **6** | **6** |
| *Chaetodon pelewensis* | CHPE | - | - | 7 | 7 | **7** | **7** |
| *Chlorurus spilurus* | CHSP | - | - | 5 | 5 | **5** | **5** |
| Category sum | | 6 | 6 | 12 | 11 | **18** | **18** |
| **Grazer/detritivores** | | | | | | | |
| *Ctenochaetus flavicauda* | CTFL | - | - | 7 | 7 | **7** | **7** |
| *Ctenochaetus striatus* | CTST | 6 | 6 | 5 | 5 | **11** | **11** |
| *Zebrosoma scopas* | ZESC | 5 | 4 | 5 | 5 | **10** | **9** |
| Category sum | | 11 | 11 | 17 | 17 | **28** | **28** |
| **Coral** | | | | | | | |
| *Acropora hyacinthus* | ACR | 6 | 4 | 6 | 2 | **12** | **6** |
| *Pocillopora* spp. | POC | 6 | 1 | 6 | 5 | **12** | **6** |
| *Porites lobata* spp. | POR | 6 | 3 | 6 | 4 | **12** | **7** |
| Category sum | | 18 | 8 | 18 | 11 | **36** | **19** |
| **Algae** | | | | | | | |
| *Asparagopsis* sp. | ASP | - | - | 6 | 5 | **6** | **5** |
| *Dictyota* sp. | DIC | 6 | 6 | - | - | **6** | **6** |
| *Lobophora* sp. | LOB | - | - | 6 | 6 | **6** | **6** |
| *Sargassum* sp. | SAR | 6 | 6 | - | - | **6** | **6** |
| *Turbinaria* sp. | TURB | 6 | 6 | 6 | 6 | **12** | **12** |
| Turf (mixed) | TURF | 6 | 6 | 5 | 5 | **11** | **11** |
| Category sum | | 24 | 24 | 23 | 22 | **47** | **46** |
| **Sediment and water** | | | | | | | |
| Sediment | SED | 5 | 5 | 5 | 4 | **10** | **9** |
| Water | WAT | 5 | 5 | 5 | 4 | **10** | **9** |
| Category sum | | 10 | 10 | 10 | 8 | **20** | **18** |

**Table S2** Pairwise comparisons of log-transformed lesion sizes (expressed as the estimated number of dead polyps) between experimental treatments. The control treatment was not included because none of the replicates had lesions. Pairwise comparisons were conducted using the package emmeans (v. 1.7.2) and we controlled for multiple comparisons using a Tukey adjustment for multiple comparisons. Significant p-values are bolded. Overall test results: Df= 3, F=4.38, p=0.02. FC: Fresh corallivore feces; FG: Fresh grazer feces; SC: Sterile corallivore feces; SG: Sterile grazer feces.

| contrast | estimate | SE | df | t.ratio | p.value |
|---|---|---|---|---|---|
| FC - FG | -1.13 | 0.425 | 16 | -2.664 | 0.075 |
| FC - SC | -0.22 | 0.491 | 16 | -0.454 | 0.968 |
| FC - SG | 0.20 | 0.491 | 16 | 0.402 | 0.977 |
| FG - SC | 0.90 | 0.425 | 16 | 2.119 | 0.189 |
| FG - SG | 1.32 | 0.425 | 16 | 3.109 | **0.031** |
| SC - SG | 0.42 | 0.491 | 16 | 0.857 | 0.827 |

**Table S3** Results from pairwise PERMANOVA tests (using Bonferroni correction for multiple comparisons) between environmental pools (algae, coral, facultative corallivore feces, grazer feces, obligate corallivore feces, and sediment and water.) Significant p-values are bolded. Because group dispersion was not homogeneous, and the sampling design was not balanced, we randomly subsampled 18 samples in each environmental pool. Samples used: Obligate corallivores: 20AMSC1, 20CHLU1, 20AMSC2, 20AMSC4, 20CHLU2, 20CHOR7, 20CHLU3, 20CHLU5, 20CHLU6, 20CHOR1, 20CHOR3, 20CHOR9, 20CHOR11, 20CHOR12, 20CHOR13, 20CHOR14, 20CHRE1, 20CHRE5; Facultative corallivores: 20CHCI5, 20CHCI6, 20CHCI1, 20CHCI2, 20CHCI3, 20CHCI4, 20CHPE3, 20CHPE4, 20CHPE5, 20CHPE1, 20CHPE2, 20CHPE6, 20CHSP1, 20CHSP2, 20CHPE7, 20CHSP3, 20CHSP4, 20CHSP5; Algae: 20Turf_pile_3, 20Turf_pile_1, 20Turf_pile_2, 20Asp_pile_1, 20Turb_pile_5, 20Lob_pile_1, 20Asp_pile_4, 20Turb_pile_3, 20Turb_pile_4, 20Turb_bak_6, 20Lob_pile_3, 20Lob_pile_4, 20Turb_bak_2, 20Turf_bak_6, 20Dict_bak_6, 20Turf_bak_2, 20Sarg_bak_4, 20Sarg_bak_5; Sediment and water: 20BakSed1, 20BakSed2, 20ForeSed1, 20ForeSed2, 20ForeSed3, 20BakSed3, 20BakSed4, 20BakSed5, 20BakWat1, 20BakWat2, 20BakWat3, 20BakWat4, 20BakWat5, 20ForeSed4, 20ForeWat1, 20ForeWat2, 20ForeWat3, 20ForeWat5; Corals: 20PocCT65B, 20PocCT15F, 20PocCt17F, 20PocCT33F, 20PocCT49F, 20PorCT68B, 20PocCT149F, 20PorCT22F, 20PorCT25F, 20PorCT136B, 20PorCT139B, 20PorCT155F, 20AcrCT64B, 20AcrCT134B, 20AcrCT94B, 20ACRBF, 20ACRCF, 20AcrCT53B; Grazer feces: 20CTFL7, 20CTST1, 20CTST2, 20CTFL1, 20CTFL3, 20CTFL5, 20CTST3, 20ZESC1, 20ZESC2, 20ZESC3, 20CTST7, 20CTST8, 20CTST9, 20CTST10, 20ZESC6, 20ZESC7, 20ZESC8, 20ZESC9.

| Comparison | | | df | Sum q | F | $R^2$ | p-value | Adjusted p-value |
|---|---|---|---|---|---|---|---|---|
| Obligate corallivore feces | - | Facultative corallivore feces | 1 | 0.941 | 2.327 | 0.064 | 0.002 | **0.036** |
| Obligate corallivore feces | - | Algae | 1 | 1.931 | 4.647 | 0.120 | 0.000 | **0.002** |
| Obligate corallivore feces | - | Sediment and water | 1 | 1.912 | 4.599 | 0.119 | 0.000 | **0.002** |
| Obligate corallivore feces | - | Coral | 1 | 1.887 | 4.775 | 0.123 | 0.000 | **0.002** |
| Obligate corallivore feces | - | Grazer feces | 1 | 2.123 | 5.369 | 0.136 | 0.000 | **0.002** |
| Facultative corallivore feces | - | Algae | 1 | 1.417 | 3.184 | 0.086 | 0.000 | **0.002** |
| Facultative corallivore feces | - | Sediment and water | 1 | 1.391 | 3.126 | 0.084 | 0.000 | **0.002** |
| Facultative corallivore feces | - | Coral | 1 | 1.496 | 3.524 | 0.094 | 0.000 | **0.002** |
| Facultative corallivore feces | - | Grazer feces | 1 | 1.382 | 3.253 | 0.087 | 0.000 | **0.002** |
| Algae | - | Sediment and water | 1 | 1.164 | 2.552 | 0.070 | 0.000 | **0.002** |
| Algae | - | Coral | 1 | 1.547 | 3.552 | 0.095 | 0.000 | **0.002** |
| Algae | - | Grazer feces | 1 | 1.550 | 3.556 | 0.095 | 0.000 | **0.002** |
| Sediment and water | - | Coral | 1 | 1.566 | 3.595 | 0.096 | 0.000 | **0.002** |
| Sediment and water | - | Grazer feces | 1 | 1.547 | 3.550 | 0.095 | 0.000 | **0.002** |
| Coral | - | Grazer feces | 1 | 1.912 | 4.602 | 0.119 | 0.000 | **0.002** |

**Table S4** Results from pairwise PERMANOVA tests between bacterial community compositions in individual species or sample types. Overall test results R2=0.409, F=2.903, p<0.001. Significant p-values are bolded. Because group dispersion was not homogeneous, and the sampling design was not balanced, we randomly subsampled 5 samples in each species/sample type. Samples used: Obligate corallivores: AMSC: 20AMSC1, 20AMSC2, 20AMSC3, 20AMSC4, 20AMSC5; CHOR: 20CHOR5, 20CHOR8, 20CHOR10, 20CHOR11, 20CHOR14; CHRE: 20CHRE8, 20CHRE1, 20CHRE3, 20CHRE4, 20CHRE7; CHLU: 20CHLU1, 20CHLU2, 20CHLU3, 20CHLU4, 20CHLU5. Facultative corallivores: CHPE: 20CHPE3, 20CHPE5, 20CHPE1, 20CHPE2, 20CHPE7; CHCI: 20CHCI5, 20CHCI6, 20CHCI1, 20CHCI3, 20CHCI4; CHSP: 20CHSP1, 20CHSP2, 20CHSP3, 20CHSP4, 20CHSP5. Grazer/detritivores: CTFL: 20CTFL1, 20CTFL2, 20CTFL3, 20CTFL4, 20CTFL5; CTST: 20CTST2, 20CTST3, 20CTST4, 20CTST6, 20CTST9; ZESC: 20ZESC1, 20ZESC3, 20ZESC5, 20ZESC8, 20ZESC9. Corals: ACR: 20AcrCT64B, 20AcrCT134B, 20AcrCT94B, 20ACRBF, 20ACRCF; POC: 20PocCT65B, 20PocCT15F, 20PocCT33F, 20PocCT49F, 20PocCT149F; POR: 20PorCT22F, 20PorCT136B, 20PorCT139B, 20PorCT146F, 20PorCT155F. Algae: Asp: 20Asp_pile_1, 20Asp_pile_2, 20Asp_pile_3, 20Asp_pile_4, 20Asp_pile_5; Dict: 20Dict_bak_1, 20Dict_bak_3, 20Dict_bak_4, 20Dict_bak_5, 20Dict_bak_6; Lob: 20Lob_pile_1, 20Lob_pile_2, 20Lob_pile_4, 20Lob_pile_5, 20Lob_pile_6; Sarg: 20Sarg_bak_1, 20Sarg_bak_2, 20Sarg_bak_3, 20Sarg_bak_5, 20Sarg_bak_6; Turb: 20Turb_pile_1, 20Turb_pile_4, 20Turb_bak_1, 20Turb_bak_3, 20Turb_bak_4; Turf: 20Turf_pile_3, 20Turf_pile_2, 20Turf_bak_6, 20Turf_bak_1, 20Turf_bak_3; Sediment and water: Sed: 20BakSed2, 20ForeSed1, 20ForeSed3, 20BakSed4, 20ForeSed4; WAT: 20BakWat4, 20BakWat5, 20ForeWat1, 20ForeWat2, 20ForeWat5.

| Comparison | | Df | Sum Sq | F | $R^2$ | p-value | Adjusted p-value |
|---|---|---|---|---|---|---|---|
| *Amanses scopas* | - *Chaetodon citrinellus* | 1 | 0.933 | 2.666 | 0.250 | 0.015 | **0.019** |
| *Amanses scopas* | - *Pocillopora* sp. | 1 | 1.391 | 5.849 | 0.422 | 0.007 | **0.013** |
| *Amanses scopas* | - *Chaetodon lunulatus* | 1 | 0.938 | 2.543 | 0.241 | 0.018 | **0.021** |
| *Amanses scopas* | - *Chaetodon ornatissimus* | 1 | 1.028 | 2.934 | 0.268 | 0.009 | **0.013** |
| *Amanses scopas* | - *Porites lobata* sp. | 1 | 0.934 | 2.445 | 0.234 | 0.016 | **0.019** |
| *Amanses scopas* | - *Sargassum* sp. | 1 | 0.759 | 1.745 | 0.179 | 0.008 | **0.013** |
| *Amanses scopas* | - *Chaetodon pelewensis* | 1 | 1.111 | 3.437 | 0.301 | 0.008 | **0.013** |
| *Amanses scopas* | - Water | 1 | 1.117 | 3.238 | 0.288 | 0.008 | **0.013** |
| *Amanses scopas* | - *Chaetodon reticulatus* | 1 | 1.135 | 3.560 | 0.308 | 0.008 | **0.013** |
| *Amanses scopas* | - Turf algae | 1 | 0.754 | 1.730 | 0.178 | 0.007 | **0.013** |
| *Amanses scopas* | - *Chlorurus spilurus* | 1 | 0.758 | 1.944 | 0.195 | 0.037 | **0.041** |
| *Amanses scopas* | - *Asparagopsis* sp. | 1 | 1.270 | 4.133 | 0.341 | 0.008 | **0.013** |
| *Amanses scopas* | - *Ctenochaetus flavicauda* | 1 | 1.173 | 3.902 | 0.328 | 0.007 | **0.013** |
| *Amanses scopas* | - *Sargassum* sp. | 1 | 1.149 | 3.408 | 0.299 | 0.006 | **0.013** |
| *Amanses scopas* | - *Ctenochaetus striatus* | 1 | 0.803 | 2.116 | 0.209 | 0.035 | **0.039** |
| *Amanses scopas* | - *Lobophora* sp. | 1 | 0.907 | 2.278 | 0.222 | 0.006 | **0.013** |
| *Amanses scopas* | - *Zebrasoma scopas* | 1 | 0.919 | 2.428 | 0.233 | 0.008 | **0.013** |
| *Amanses scopas* | - *Dictyota* sp. | 1 | 0.847 | 2.060 | 0.205 | 0.010 | **0.013** |
| *Amanses scopas* | - *Sargassum* sp. | 1 | 0.821 | 1.960 | 0.197 | 0.010 | **0.013** |
| *Amanses scopas* | - *Acropora hyacinthus* | 1 | 0.824 | 1.971 | 0.198 | 0.008 | **0.013** |
| *Chaetodon citrinellus* | - *Pocillopora* sp. | 1 | 1.655 | 7.928 | 0.498 | 0.009 | **0.013** |

| | | | | | | | |
|---|---|---|---|---|---|---|---|
| *Chaetodon citrinellus* | - *Chaetodon lunulatus* | 1 | 0.874 | 2.570 | 0.243 | 0.015 | **0.019** |
| *Chaetodon citrinellus* | - *Chaetodon ornatissimus* | 1 | 1.159 | 3.608 | 0.311 | 0.010 | **0.013** |
| *Chaetodon citrinellus* | - *Porites lobata* sp. | 1 | 1.078 | 3.052 | 0.276 | 0.006 | **0.013** |
| *Chaetodon citrinellus* | - *Sargassum* sp. | 1 | 0.877 | 2.162 | 0.213 | 0.008 | **0.013** |
| *Chaetodon citrinellus* | - *Chaetodon pelewensis* | 1 | 1.262 | 4.287 | 0.349 | 0.009 | **0.013** |
| *Chaetodon citrinellus* | - Water | 1 | 1.236 | 3.910 | 0.328 | 0.009 | **0.013** |
| *Chaetodon citrinellus* | - *Chaetodon reticulatus* | 1 | 1.289 | 4.447 | 0.357 | 0.008 | **0.013** |
| *Chaetodon citrinellus* | - Turf algae | 1 | 0.873 | 2.147 | 0.212 | 0.007 | **0.013** |
| *Chaetodon citrinellus* | - *Chlorurus spilurus* | 1 | 1.040 | 2.880 | 0.265 | 0.009 | **0.013** |
| *Chaetodon citrinellus* | - *Asparagopsis* sp. | 1 | 1.387 | 4.986 | 0.384 | 0.009 | **0.013** |
| *Chaetodon citrinellus* | - *Ctenochaetus flavicauda* | 1 | 1.391 | 5.122 | 0.390 | 0.009 | **0.013** |
| *Chaetodon citrinellus* | - *Sargassum* sp. | 1 | 1.267 | 4.111 | 0.339 | 0.009 | **0.013** |
| *Chaetodon citrinellus* | - *Ctenochaetus striatus* | 1 | 1.090 | 3.111 | 0.280 | 0.007 | **0.013** |
| *Chaetodon citrinellus* | - *Lobophora* sp. | 1 | 1.024 | 2.776 | 0.258 | 0.008 | **0.013** |
| *Chaetodon citrinellus* | - *Zebrasoma scopas* | 1 | 1.077 | 3.083 | 0.278 | 0.007 | **0.013** |
| *Chaetodon citrinellus* | - *Dictyota* sp. | 1 | 0.971 | 2.542 | 0.241 | 0.009 | **0.013** |
| *Chaetodon citrinellus* | - *Sargassum* sp. | 1 | 0.941 | 2.415 | 0.232 | 0.008 | **0.013** |
| *Chaetodon citrinellus* | - *Acropora hyacinthus* | 1 | 0.944 | 2.429 | 0.233 | 0.008 | **0.013** |
| *Pocillopora* sp. | - *Chaetodon lunulatus* | 1 | 1.577 | 6.920 | 0.464 | 0.010 | **0.013** |
| *Pocillopora* sp. | - *Chaetodon ornatissimus* | 1 | 1.648 | 7.880 | 0.496 | 0.008 | **0.013** |
| *Pocillopora* sp. | - *Porites lobata* sp. | 1 | 1.152 | 4.780 | 0.374 | 0.018 | **0.021** |
| *Pocillopora* sp. | - *Sargassum* sp. | 1 | 1.322 | 4.502 | 0.360 | 0.009 | **0.013** |
| *Pocillopora* sp. | - *Chaetodon pelewensis* | 1 | 1.660 | 9.106 | 0.532 | 0.007 | **0.013** |
| *Pocillopora* sp. | - Water | 1 | 1.677 | 8.219 | 0.507 | 0.010 | **0.013** |
| *Pocillopora* sp. | - *Chaetodon reticulatus* | 1 | 1.700 | 9.556 | 0.544 | 0.010 | **0.013** |
| *Pocillopora* sp. | - Turf algae | 1 | 1.322 | 4.486 | 0.359 | 0.007 | **0.013** |
| *Pocillopora* sp. | - *Chlorurus spilurus* | 1 | 1.471 | 5.909 | 0.425 | 0.009 | **0.013** |
| *Pocillopora* sp. | - *Asparagopsis* sp. | 1 | 1.834 | 11.033 | 0.580 | 0.009 | **0.013** |
| *Pocillopora* sp. | - *Ctenochaetus flavicauda* | 1 | 1.862 | 11.663 | 0.593 | 0.008 | **0.013** |
| *Pocillopora* sp. | - *Sargassum* sp. | 1 | 1.715 | 8.740 | 0.522 | 0.008 | **0.013** |
| *Pocillopora* sp. | - *Ctenochaetus striatus* | 1 | 1.542 | 6.470 | 0.447 | 0.009 | **0.013** |
| *Pocillopora* sp. | - *Lobophora* sp. | 1 | 1.471 | 5.723 | 0.417 | 0.008 | **0.013** |
| *Pocillopora* sp. | - *Zebrasoma scopas* | 1 | 1.544 | 6.505 | 0.448 | 0.009 | **0.013** |
| *Pocillopora* sp. | - *Dictyota* sp. | 1 | 1.420 | 5.260 | 0.397 | 0.006 | **0.013** |
| *Pocillopora* sp. | - *Sargassum* sp. | 1 | 1.390 | 5.007 | 0.385 | 0.008 | **0.013** |
| *Pocillopora* sp. | - *Acropora hyacinthus* | 1 | 1.391 | 5.027 | 0.386 | 0.009 | **0.013** |
| *Chaetodon lunulatus* | - *Chaetodon ornatissimus* | 1 | 0.364 | 1.070 | 0.118 | 0.215 | 0.219 |
| *Chaetodon lunulatus* | - *Porites lobata* sp. | 1 | 0.672 | 1.806 | 0.184 | 0.016 | **0.019** |
| *Chaetodon lunulatus* | - *Sargassum* sp. | 1 | 0.789 | 1.858 | 0.188 | 0.008 | **0.013** |
| *Chaetodon lunulatus* | - *Chaetodon pelewensis* | 1 | 0.504 | 1.608 | 0.167 | 0.015 | **0.018** |
| *Chaetodon lunulatus* | - Water | 1 | 1.159 | 3.459 | 0.302 | 0.008 | **0.013** |
| *Chaetodon lunulatus* | - *Chaetodon reticulatus* | 1 | 0.441 | 1.427 | 0.151 | 0.007 | **0.013** |
| *Chaetodon lunulatus* | - Turf algae | 1 | 0.797 | 1.872 | 0.190 | 0.009 | **0.013** |
| *Chaetodon lunulatus* | - *Chlorurus spilurus* | 1 | 0.785 | 2.065 | 0.205 | 0.017 | **0.020** |
| *Chaetodon lunulatus* | - *Asparagopsis* sp. | 1 | 1.311 | 4.409 | 0.355 | 0.007 | **0.013** |
| *Chaetodon lunulatus* | - *Ctenochaetus flavicauda* | 1 | 1.307 | 4.497 | 0.360 | 0.010 | **0.013** |
| *Chaetodon lunulatus* | - *Sargassum* sp. | 1 | 1.191 | 3.638 | 0.313 | 0.008 | **0.013** |
| *Chaetodon lunulatus* | - *Ctenochaetus striatus* | 1 | 1.001 | 2.711 | 0.253 | 0.007 | **0.013** |
| *Chaetodon lunulatus* | - *Lobophora* sp. | 1 | 0.948 | 2.443 | 0.234 | 0.008 | **0.013** |
| *Chaetodon lunulatus* | - *Zebrasoma scopas* | 1 | 1.006 | 2.731 | 0.255 | 0.010 | **0.013** |
| *Chaetodon lunulatus* | - *Dictyota* sp. | 1 | 0.895 | 2.231 | 0.218 | 0.008 | **0.013** |
| *Chaetodon lunulatus* | - *Sargassum* sp. | 1 | 0.865 | 2.116 | 0.209 | 0.008 | **0.013** |
| *Chaetodon lunulatus* | - *Acropora hyacinthus* | 1 | 0.868 | 2.129 | 0.210 | 0.007 | **0.013** |
| *Chaetodon ornatissimus* | - *Porites lobata* sp. | 1 | 0.477 | 1.349 | 0.144 | 0.154 | 0.160 |
| *Chaetodon ornatissimus* | - *Sargassum* sp. | 1 | 0.858 | 2.113 | 0.209 | 0.007 | **0.013** |
| *Chaetodon ornatissimus* | - *Chaetodon pelewensis* | 1 | 0.401 | 1.361 | 0.145 | 0.098 | 0.104 |
| *Chaetodon ornatissimus* | - Water | 1 | 1.234 | 3.899 | 0.328 | 0.008 | **0.013** |
| *Chaetodon ornatissimus* | - *Chaetodon reticulatus* | 1 | 0.318 | 1.095 | 0.120 | 0.272 | 0.275 |

| | | | | | | | |
|---|---|---|---|---|---|---|---|
| *Chaetodon ornatissimus* | - Turf algae | 1 | 0.872 | 2.142 | 0.211 | 0.008 | **0.013** |
| *Chaetodon ornatissimus* | - *Chlorurus spilurus* | 1 | 0.837 | 2.318 | 0.225 | 0.034 | **0.038** |
| *Chaetodon ornatissimus* | - *Asparagopsis* sp. | 1 | 1.386 | 4.974 | 0.383 | 0.008 | **0.013** |
| *Chaetodon ornatissimus* | - *Ctenochaetus flavicauda* | 1 | 1.391 | 5.115 | 0.390 | 0.010 | **0.013** |
| *Chaetodon ornatissimus* | - *Sargassum* sp. | 1 | 1.266 | 4.101 | 0.339 | 0.009 | **0.013** |
| *Chaetodon ornatissimus* | - *Ctenochaetus striatus* | 1 | 1.084 | 3.090 | 0.279 | 0.008 | **0.013** |
| *Chaetodon ornatissimus* | - *Lobophora* sp. | 1 | 1.023 | 2.769 | 0.257 | 0.008 | **0.013** |
| *Chaetodon ornatissimus* | - *Zebrasoma scopas* | 1 | 1.084 | 3.100 | 0.279 | 0.008 | **0.013** |
| *Chaetodon ornatissimus* | - *Dictyota* sp. | 1 | 0.971 | 2.539 | 0.241 | 0.007 | **0.013** |
| *Chaetodon ornatissimus* | - *Sargassum* sp. | 1 | 0.940 | 2.411 | 0.232 | 0.010 | **0.013** |
| *Chaetodon ornatissimus* | - *Acropora hyacinthus* | 1 | 0.943 | 2.425 | 0.233 | 0.008 | **0.013** |
| *Porites lobata* sp. | - *Sargassum* sp. | 1 | 0.722 | 1.650 | 0.171 | 0.024 | **0.027** |
| *Porites lobata* sp. | - *Chaetodon pelewensis* | 1 | 0.492 | 1.505 | 0.158 | 0.089 | 0.095 |
| *Porites lobata* sp. | - Water | 1 | 1.103 | 3.166 | 0.284 | 0.008 | **0.013** |
| *Porites lobata* sp. | - *Chaetodon reticulatus* | 1 | 0.472 | 1.466 | 0.155 | 0.119 | 0.124 |
| *Porites lobata* sp. | - Turf algae | 1 | 0.744 | 1.695 | 0.175 | 0.007 | **0.013** |
| *Porites lobata* sp. | - *Chlorurus spilurus* | 1 | 0.819 | 2.084 | 0.207 | 0.025 | **0.028** |
| *Porites lobata* sp. | - *Asparagopsis* sp. | 1 | 1.252 | 4.032 | 0.335 | 0.017 | **0.020** |
| *Porites lobata* sp. | - *Ctenochaetus flavicauda* | 1 | 1.270 | 4.180 | 0.343 | 0.008 | **0.013** |
| *Porites lobata* sp. | - *Sargassum* sp. | 1 | 1.138 | 3.342 | 0.295 | 0.009 | **0.013** |
| *Porites lobata* sp. | - *Ctenochaetus striatus* | 1 | 0.961 | 2.512 | 0.239 | 0.007 | **0.013** |
| *Porites lobata* sp. | - *Lobophora* sp. | 1 | 0.885 | 2.205 | 0.216 | 0.008 | **0.013** |
| *Porites lobata* sp. | - *Zebrasoma scopas* | 1 | 0.965 | 2.527 | 0.240 | 0.009 | **0.013** |
| *Porites lobata* sp. | - *Dictyota* sp. | 1 | 0.843 | 2.036 | 0.203 | 0.007 | **0.013** |
| *Porites lobata* sp. | - *Sargassum* sp. | 1 | 0.812 | 1.924 | 0.194 | 0.008 | **0.013** |
| *Porites lobata* sp. | - *Acropora hyacinthus* | 1 | 0.805 | 1.911 | 0.193 | 0.023 | **0.027** |
| Sediment | - *Chaetodon pelewensis* | 1 | 0.956 | 2.521 | 0.240 | 0.009 | **0.013** |
| Sediment | - Water | 1 | 0.852 | 2.125 | 0.210 | 0.008 | **0.013** |
| Sediment | - *Chaetodon reticulatus* | 1 | 0.973 | 2.596 | 0.245 | 0.008 | **0.013** |
| Sediment | - Turf algae | 1 | 0.491 | 1.000 | 0.111 | 0.443 | 0.445 |
| Sediment | - *Chlorurus spilurus* | 1 | 0.702 | 1.575 | 0.164 | 0.009 | **0.013** |
| Sediment | - *Asparagopsis* sp. | 1 | 1.009 | 2.778 | 0.258 | 0.048 | 0.053 |
| Sediment | - *Ctenochaetus flavicauda* | 1 | 1.066 | 2.990 | 0.272 | 0.008 | **0.013** |
| Sediment | - *Sargassum* sp. | 1 | 0.877 | 2.232 | 0.218 | 0.008 | **0.013** |
| Sediment | - *Ctenochaetus striatus* | 1 | 0.754 | 1.732 | 0.178 | 0.008 | **0.013** |
| Sediment | - *Lobophora* sp. | 1 | 0.581 | 1.281 | 0.138 | 0.056 | 0.061 |
| Sediment | - *Zebrasoma scopas* | 1 | 0.760 | 1.750 | 0.179 | 0.008 | **0.013** |
| Sediment | - *Dictyota* sp. | 1 | 0.604 | 1.293 | 0.139 | 0.007 | **0.013** |
| Sediment | - *Sargassum* sp. | 1 | 0.593 | 1.249 | 0.135 | 0.025 | **0.028** |
| Sediment | - *Acropora hyacinthus* | 1 | 0.601 | 1.269 | 0.137 | 0.023 | **0.027** |
| *Chaetodon pelewensis* | - Water | 1 | 1.341 | 4.631 | 0.367 | 0.008 | **0.013** |
| *Chaetodon pelewensis* | - *Chaetodon reticulatus* | 1 | 0.307 | 1.167 | 0.127 | 0.199 | 0.204 |
| *Chaetodon pelewensis* | - Turf algae | 1 | 0.980 | 2.577 | 0.244 | 0.007 | **0.013** |
| *Chaetodon pelewensis* | - *Chlorurus spilurus* | 1 | 0.944 | 2.823 | 0.261 | 0.018 | **0.021** |
| *Chaetodon pelewensis* | - *Asparagopsis* sp. | 1 | 1.493 | 5.932 | 0.426 | 0.006 | **0.013** |
| *Chaetodon pelewensis* | - *Ctenochaetus flavicauda* | 1 | 1.501 | 6.125 | 0.434 | 0.010 | **0.013** |
| *Chaetodon pelewensis* | - *Sargassum* sp. | 1 | 1.373 | 4.875 | 0.379 | 0.007 | **0.013** |
| *Chaetodon pelewensis* | - *Ctenochaetus striatus* | 1 | 1.189 | 3.672 | 0.315 | 0.010 | **0.013** |
| *Chaetodon pelewensis* | - *Lobophora* sp. | 1 | 1.130 | 3.301 | 0.292 | 0.009 | **0.013** |
| *Chaetodon pelewensis* | - *Zebrasoma scopas* | 1 | 1.193 | 3.696 | 0.316 | 0.008 | **0.013** |
| *Chaetodon pelewensis* | - *Dictyota* sp. | 1 | 1.078 | 3.034 | 0.275 | 0.009 | **0.013** |
| *Chaetodon pelewensis* | - *Sargassum* sp. | 1 | 1.048 | 2.886 | 0.265 | 0.009 | **0.013** |
| *Chaetodon pelewensis* | - *Acropora hyacinthus* | 1 | 1.051 | 2.902 | 0.266 | 0.007 | **0.013** |
| Water | - *Chaetodon reticulatus* | 1 | 1.359 | 4.765 | 0.373 | 0.009 | **0.013** |
| Water | - Turf algae | 1 | 0.883 | 2.197 | 0.215 | 0.009 | **0.013** |
| Water | - *Chlorurus spilurus* | 1 | 1.073 | 3.013 | 0.274 | 0.008 | **0.013** |
| Water | - *Asparagopsis* sp. | 1 | 1.404 | 5.133 | 0.391 | 0.009 | **0.013** |
| Water | - *Ctenochaetus flavicauda* | 1 | 1.431 | 5.362 | 0.401 | 0.009 | **0.013** |

| | | | | | | | |
|---|---|---|---|---|---|---|---|
| Water | - *Sargassum* sp. | 1 | 1.281 | 4.222 | 0.345 | 0.007 | **0.013** |
| Water | - *Ctenochaetus striatus* | 1 | 1.117 | 3.234 | 0.288 | 0.007 | **0.013** |
| Water | - *Lobophora* sp. | 1 | 1.041 | 2.859 | 0.263 | 0.008 | **0.013** |
| Water | - *Zebrasoma scopas* | 1 | 1.121 | 3.254 | 0.289 | 0.008 | **0.013** |
| Water | - *Dictyota* sp. | 1 | 0.990 | 2.624 | 0.247 | 0.008 | **0.013** |
| Water | - *Sargassum* sp. | 1 | 0.959 | 2.493 | 0.238 | 0.008 | **0.013** |
| Water | - *Acropora hyacinthus* | 1 | 0.954 | 2.485 | 0.237 | 0.007 | **0.013** |
| *Chaetodon reticulatus* | - Turf algae | 1 | 0.997 | 2.654 | 0.249 | 0.009 | **0.013** |
| *Chaetodon reticulatus* | - *Chlorurus spilurus* | 1 | 0.918 | 2.781 | 0.258 | 0.023 | **0.027** |
| *Chaetodon reticulatus* | - *Asparagopsis* sp. | 1 | 1.511 | 6.110 | 0.433 | 0.006 | **0.013** |
| *Chaetodon reticulatus* | - *Ctenochaetus flavicauda* | 1 | 1.509 | 6.270 | 0.439 | 0.007 | **0.013** |
| *Chaetodon reticulatus* | - *Sargassum* sp. | 1 | 1.391 | 5.016 | 0.385 | 0.009 | **0.013** |
| *Chaetodon reticulatus* | - *Ctenochaetus striatus* | 1 | 1.205 | 3.773 | 0.320 | 0.008 | **0.013** |
| *Chaetodon reticulatus* | - *Lobophora* sp. | 1 | 1.148 | 3.396 | 0.298 | 0.008 | **0.013** |
| *Chaetodon reticulatus* | - *Zebrasoma scopas* | 1 | 1.209 | 3.797 | 0.322 | 0.008 | **0.013** |
| *Chaetodon reticulatus* | - *Dictyota* sp. | 1 | 1.096 | 3.122 | 0.281 | 0.008 | **0.013** |
| *Chaetodon reticulatus* | - *Sargassum* sp. | 1 | 1.065 | 2.970 | 0.271 | 0.007 | **0.013** |
| *Chaetodon reticulatus* | - *Acropora hyacinthus* | 1 | 1.069 | 2.987 | 0.272 | 0.007 | **0.013** |
| Turf algae | - *Chlorurus spilurus* | 1 | 0.700 | 1.567 | 0.164 | 0.009 | **0.013** |
| Turf algae | - *Asparagopsis* sp. | 1 | 1.030 | 2.828 | 0.261 | 0.016 | **0.019** |
| Turf algae | - *Ctenochaetus flavicauda* | 1 | 1.058 | 2.959 | 0.270 | 0.008 | **0.013** |
| Turf algae | - *Sargassum* sp. | 1 | 0.901 | 2.288 | 0.222 | 0.008 | **0.013** |
| Turf algae | - *Ctenochaetus striatus* | 1 | 0.743 | 1.705 | 0.176 | 0.008 | **0.013** |
| Turf algae | - *Lobophora* sp. | 1 | 0.653 | 1.436 | 0.152 | 0.024 | **0.028** |
| Turf algae | - *Zebrasoma scopas* | 1 | 0.758 | 1.741 | 0.179 | 0.010 | **0.013** |
| Turf algae | - *Dictyota* sp. | 1 | 0.594 | 1.270 | 0.137 | 0.007 | **0.013** |
| Turf algae | - *Sargassum* sp. | 1 | 0.552 | 1.161 | 0.127 | 0.082 | 0.088 |
| Turf algae | - *Acropora hyacinthus* | 1 | 0.586 | 1.236 | 0.134 | 0.070 | 0.074 |
| *Chlorurus spilurus* | - *Asparagopsis* sp. | 1 | 1.223 | 3.841 | 0.324 | 0.008 | **0.013** |
| *Chlorurus spilurus* | - *Ctenochaetus flavicauda* | 1 | 0.882 | 2.829 | 0.261 | 0.008 | **0.013** |
| *Chlorurus spilurus* | - *Sargassum* sp. | 1 | 1.098 | 3.153 | 0.283 | 0.008 | **0.013** |
| *Chlorurus spilurus* | - *Ctenochaetus striatus* | 1 | 0.484 | 1.241 | 0.134 | 0.179 | 0.186 |
| *Chlorurus spilurus* | - *Lobophora* sp. | 1 | 0.855 | 2.089 | 0.207 | 0.008 | **0.013** |
| *Chlorurus spilurus* | - *Zebrasoma scopas* | 1 | 0.764 | 1.962 | 0.197 | 0.022 | **0.027** |
| *Chlorurus spilurus* | - *Dictyota* sp. | 1 | 0.803 | 1.903 | 0.192 | 0.008 | **0.013** |
| *Chlorurus spilurus* | - *Sargassum* sp. | 1 | 0.771 | 1.795 | 0.183 | 0.008 | **0.013** |
| *Chlorurus spilurus* | - *Acropora hyacinthus* | 1 | 0.782 | 1.822 | 0.186 | 0.007 | **0.013** |
| *Asparagopsis* sp. | - *Ctenochaetus flavicauda* | 1 | 1.581 | 6.905 | 0.463 | 0.008 | **0.013** |
| *Asparagopsis* sp. | - *Sargassum* sp. | 1 | 1.428 | 5.376 | 0.402 | 0.009 | **0.013** |
| *Asparagopsis* sp. | - *Ctenochaetus striatus* | 1 | 1.262 | 4.101 | 0.339 | 0.007 | **0.013** |
| *Asparagopsis* sp. | - *Lobophora* sp. | 1 | 1.001 | 3.066 | 0.277 | 0.050 | 0.055 |
| *Asparagopsis* sp. | - *Zebrasoma scopas* | 1 | 1.272 | 4.147 | 0.341 | 0.009 | **0.013** |
| *Asparagopsis* sp. | - *Dictyota* sp. | 1 | 1.120 | 3.302 | 0.292 | 0.010 | **0.013** |
| *Asparagopsis* sp. | - *Sargassum* sp. | 1 | 1.104 | 3.181 | 0.284 | 0.025 | **0.028** |
| *Asparagopsis* sp. | - *Acropora hyacinthus* | 1 | 1.111 | 3.210 | 0.286 | 0.008 | **0.013** |
| *Ctenochaetus flavicauda* | - *Sargassum* sp. | 1 | 1.453 | 5.608 | 0.412 | 0.007 | **0.013** |
| *Ctenochaetus flavicauda* | - *Ctenochaetus striatus* | 1 | 0.979 | 3.250 | 0.289 | 0.008 | **0.013** |
| *Ctenochaetus flavicauda* | - *Lobophora* sp. | 1 | 1.215 | 3.800 | 0.322 | 0.008 | **0.013** |
| *Ctenochaetus flavicauda* | - *Zebrasoma scopas* | 1 | 1.018 | 3.391 | 0.298 | 0.007 | **0.013** |
| *Ctenochaetus flavicauda* | - *Dictyota* sp. | 1 | 1.141 | 3.431 | 0.300 | 0.010 | **0.013** |
| *Ctenochaetus flavicauda* | - *Sargassum* sp. | 1 | 1.127 | 3.309 | 0.293 | 0.009 | **0.013** |
| *Ctenochaetus flavicauda* | - *Acropora hyacinthus* | 1 | 1.139 | 3.354 | 0.295 | 0.011 | **0.013** |
| *Turbinaria* sp. | - *Ctenochaetus striatus* | 1 | 1.134 | 3.359 | 0.296 | 0.008 | **0.013** |
| *Turbinaria* sp. | - *Lobophora* sp. | 1 | 0.743 | 2.085 | 0.207 | 0.050 | 0.055 |
| *Turbinaria* sp. | - *Zebrasoma scopas* | 1 | 1.144 | 3.396 | 0.298 | 0.008 | **0.013** |
| *Turbinaria* sp. | - *Dictyota* sp. | 1 | 0.834 | 2.258 | 0.220 | 0.010 | **0.013** |
| *Turbinaria* sp. | - *Sargassum* sp. | 1 | 0.924 | 2.452 | 0.235 | 0.007 | **0.013** |
| *Turbinaria* sp. | - *Acropora hyacinthus* | 1 | 0.940 | 2.499 | 0.238 | 0.008 | **0.013** |

| | | | | | | | |
|---|---|---|---|---|---|---|---|
| *Ctenochaetus striatus* | - *Lobophora* sp. | 1 | 0.884 | 2.219 | 0.217 | 0.007 | **0.013** |
| *Ctenochaetus striatus* | - *Zebrasoma scopas* | 1 | 0.577 | 1.524 | 0.160 | 0.197 | 0.203 |
| *Ctenochaetus striatus* | - *Dictyota* sp. | 1 | 0.839 | 2.040 | 0.203 | 0.007 | **0.013** |
| *Ctenochaetus striatus* | - *Sargassum* sp. | 1 | 0.813 | 1.940 | 0.195 | 0.008 | **0.013** |
| *Ctenochaetus striatus* | - *Acropora hyacinthus* | 1 | 0.826 | 1.975 | 0.198 | 0.009 | **0.013** |
| Lobophora sp. | - *Zebrasoma scopas* | 1 | 0.907 | 2.282 | 0.222 | 0.009 | **0.013** |
| *Lobophora* sp. | - *Dictyota* sp. | 1 | 0.639 | 1.485 | 0.157 | 0.108 | 0.113 |
| *Lobophora* sp. | - *Sargassum* sp. | 1 | 0.660 | 1.509 | 0.159 | 0.047 | 0.052 |
| *Lobophora* sp. | - *Acropora hyacinthus* | 1 | 0.699 | 1.600 | 0.167 | 0.048 | 0.053 |
| *Zebrasoma scopas* | - *Dictyota* sp. | 1 | 0.851 | 2.074 | 0.206 | 0.009 | **0.013** |
| *Zebrasoma scopas* | - *Sargassum* sp. | 1 | 0.826 | 1.975 | 0.198 | 0.007 | **0.013** |
| *Zebrasoma scopas* | - *Acropora hyacinthus* | 1 | 0.831 | 1.991 | 0.199 | 0.008 | **0.013** |
| *Dictyota* sp. | - *Sargassum* sp. | 1 | 0.436 | 0.968 | 0.108 | 0.685 | 0.685 |
| *Dictyota* sp. | - *Acropora hyacinthus* | 1 | 0.612 | 1.361 | 0.145 | 0.062 | 0.067 |
| *Sargassum* sp. | - *Acropora hyacinthus* | 1 | 0.518 | 1.133 | 0.124 | 0.235 | 0.239 |

**Table S5** Number of bacterial ASVs associated with environmental pools (resource species corals or algae, or sediment or water) that were also identified in consumer feces: for example, feces of the obligate corallivore CHRE (*Chaetodon reticulatus*) contained 225 of 254 ASVs that were identified in colonies of POR (*Porites lobata* species complex). Darker green shading indicates higher percentages; shading was scaled for each row (i.e., each consumer species) separately. For this analysis, all sequencing libraries were rarefied to 1,037 reads per sample, and all samples per species or environmental reservoirs were pooled. "N# bacterial ASVs" indicate the total number of unique ASVS in each environmental reservoir or fish species after rarefaction. See Figure for the percentages of shared ASVs. See table S1 for species names.

| | | Environmental reservoir and N# bacterial ASVs | | | | | | | | | | |
|---|---|---|---|---|---|---|---|---|---|---|---|---|
| Fish species | N# Bacterial ASVs | Corals | | | Algae | | | | | | Sediment and water | |
| | | ACR | POC | POR | Asp | Dict | Lob | Sarg | Turb | Turf | Sed | Wat |
| | | 130 | 119 | 254 | 254 | 385 | 450 | 233 | 548 | 425 | 261 | 343 |
| *Obligate corallivores* | | | | | | | | | | | | |
| CHRE | 1383 | 0 | 83 | 225 | 2 | 1 | 1 | 0 | 0 | 0 | 13 | 8 |
| CHOR | 1845 | 0 | 71 | 221 | 2 | 0 | 1 | 0 | 0 | 3 | 18 | 11 |
| CHLU | 1424 | 0 | 18 | 159 | 1 | 6 | 0 | 4 | 0 | 2 | 10 | 20 |
| AMSC | 702 | 4 | 90 | 75 | 2 | 18 | 1 | 8 | 7 | 10 | 28 | 74 |
| *Facultative corallivores* | | | | | | | | | | | | |
| CHPE | 1294 | 2 | 76 | 218 | 26 | 23 | 33 | 15 | 11 | 53 | 27 | 14 |
| CHCI | 444 | 1 | 6 | 11 | 0 | 7 | 1 | 2 | 1 | 3 | 4 | 22 |
| CHSP | 1213 | 2 | 33 | 149 | 11 | 25 | 20 | 17 | 13 | 37 | 43 | 51 |
| *Grazer/detritivores* | | | | | | | | | | | | |
| CTFL | 816 | 3 | 2 | 11 | 25 | 40 | 36 | 24 | 27 | 68 | 50 | 88 |
| CTST | 1164 | 6 | 6 | 23 | 30 | 60 | 57 | 45 | 39 | 89 | 65 | 91 |
| ZESC | 874 | 1 | 4 | 14 | 5 | 20 | 8 | 10 | 15 | 23 | 26 | 37 |

**Table S6** Results of pairwise comparisons using Dunn tests with Benjamini-Hochberg correction of Endozoicomonadaceae relative read abundances between different sample types (algae, sediment and water, coral, grazer feces, facultative corallivore feces, and obligate corallivore feces). Overall Kruskal-Wallis test results: $X^2$=94.5, df=5, p<0.001.

| Comparison | | | Z statistic | p-value |
|---|---|---|---|---|
| Algae | - | Coral | -6.150 | **0.000** |
| Algae | - | Sediment and water | -0.720 | 0.295 |
| Coral | - | Sediment and water | 4.490 | **0.000** |
| Algae | - | Facultative corallivore feces | -6.434 | **0.000** |
| Coral | - | Facultative corallivore feces | -0.340 | 0.424 |
| Sediment and water | - | Facultative corallivore feces | -4.766 | **0.000** |
| Algae | - | Grazer feces | -4.230 | **0.000** |
| Coral | - | Grazer feces | 2.231 | **0.018** |
| Sediment and water | - | Grazer feces | -2.693 | **0.006** |
| Facultative corallivore feces | - | Grazer feces | 2.565 | **0.008** |
| Algae | - | Obligate corallivore feces | -7.563 | **0.000** |
| Coral | - | Obligate corallivore feces | -0.221 | 0.442 |
| Sediment and water | - | Obligate corallivore feces | -5.230 | **0.000** |
| Facultative corallivore feces | - | Obligate corallivore feces | 0.162 | 0.436 |
| Grazer feces | - | Obligate corallivore feces | -2.810 | **0.005** |

**Table S7** Results of Kruskal-Wallis tests for differences in Endozoicomonadaceae relative read abundances between species within each environmental pool (coral, algae, sediment and water, grazer feces, facultative corallivore feces, and obligate corallivore feces), as well as pairwise comparisons (Dunn tests) between species or sample types within each environmental pool. N.A.: No comparisons; N.S.: Kruskal-Wallis test not significant.

| Comparison | | | Dunn test results | |
|---|---|---|---|---|
| | | | *Z statistic* | *p-value* |
| *Algae (Kruskal-Wallis test results: $X^2$=12.248, df=5,* ***p=0.032****)* | | | | |
| *Asparagopsis* sp. | - | *Dictyota* sp. | 0.295 | 0.412 |
| *Asparagopsis* sp. | - | *Lobophora* sp. | 1.769 | 0.192 |
| *Dictyota* sp. | - | *Lobophora* sp. | 1.546 | 0.153 |
| *Asparagopsis* sp. | - | *Sargassum* sp. | 0.295 | 0.443 |
| *Dictyota* sp. | - | *Sargassum* sp. | 0.000 | 0.500 |
| *Lobophora* sp. | - | *Sargassum* sp. | -1.546 | 0.183 |
| *Asparagopsis* sp. | - | *Turbinaria* sp. | -0.524 | 0.410 |
| *Dictyota* sp. | - | *Turbinaria* sp. | -0.914 | 0.270 |
| *Lobophora* sp. | - | *Turbinaria* sp. | -2.700 | **0.026** |
| *Sargassum* sp. | - | *Turbinaria* sp. | -0.914 | 0.300 |
| *Asparagopsis* sp. | - | Turf (mixed) | 1.685 | 0.172 |
| *Dictyota* sp. | - | Turf (mixed) | 1.439 | 0.141 |
| *Lobophora* sp. | - | Turf (mixed) | -0.320 | 0.468 |
| *Sargassum* sp. | - | Turf (mixed) | 1.439 | 0.161 |
| *Turbinaria* sp. | - | Turf (mixed) | 2.845 | **0.033** |
| *Coral (Kruskal-Wallis test results: $X^2$=12.7, df = 2,* ***p=0.002****)* | | | | |
| *Acropora hyacinthus* | - | *Pocillopora* spp. | -3.520 | **0.001** |
| *Acropora hyacinthus* | - | *Porites lobata* spp. | -2.193 | **0.021** |
| *Pocillopora* spp. | - | *Porites lobata* spp. | 1.512 | 0.065 |
| *Facultative corallivore feces (Kruskal-Wallis test results: $X^2$=4.318, df=2, p=0.115)* | | | | |
| N.S. | | | | |
| *Grazer feces (Kruskal-Wallis test results: $X^2$=0.588, df=2, p=0.750)* | | | | |
| N.S. | | | | |
| *Obligate corallivore feces (Kruskal-Wallis test results: $X^2$=8.892, df=3,* ***p=0.031****)* | | | | |
| *Amanses scopas* | - | *Chaetodon lunulatus* | -1.456 | 0.146 |
| *Amanses scopas* | - | *Chaetodon ornatissimus* | -2.532 | **0.017** |
| *Chaetodon lunulatus* | - | *Chaetodon ornatissimus* | -0.897 | 0.222 |
| *Amanses scopas* | - | *Chaetodon reticulatus* | -2.764 | **0.017** |
| *Chaetodon lunulatus* | - | *Chaetodon reticulatus* | -1.325 | 0.139 |
| *Chaetodon ornatissimus* | - | *Chaetodon reticulatus* | -0.647 | 0.259 |
| *Sediment and water (Kruskal-Wallis test results: $X^2$=0.007, df=1, p-value=0.932)* | | | | |
| N.S. | | | | |

### Supplementary Figures

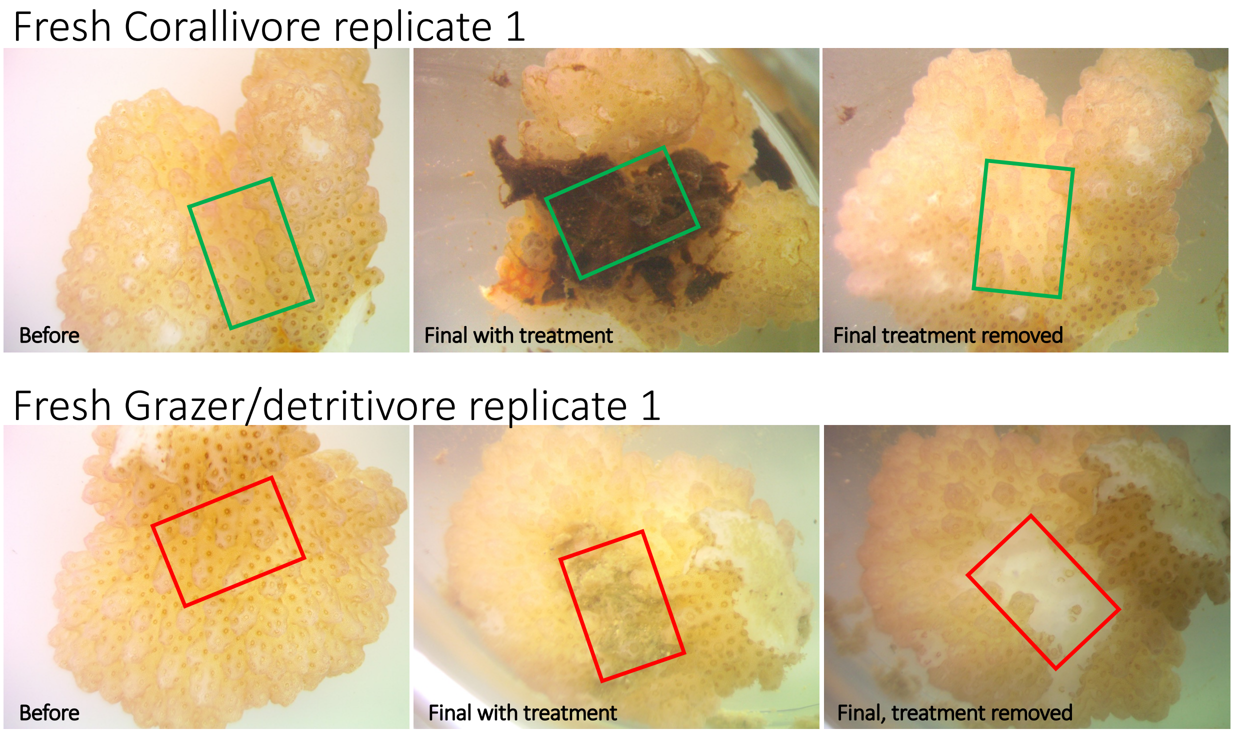


**Figure S1** Example pictures of coral fragments with fresh corallivore feces or grazer/detritivore feces treatments. Grazer/detritivore feces caused lesions or mortality in all replicates (N=11/11), whereas fresh corallivore feces caused lesions in only 45% (N=5/11) of fragments.

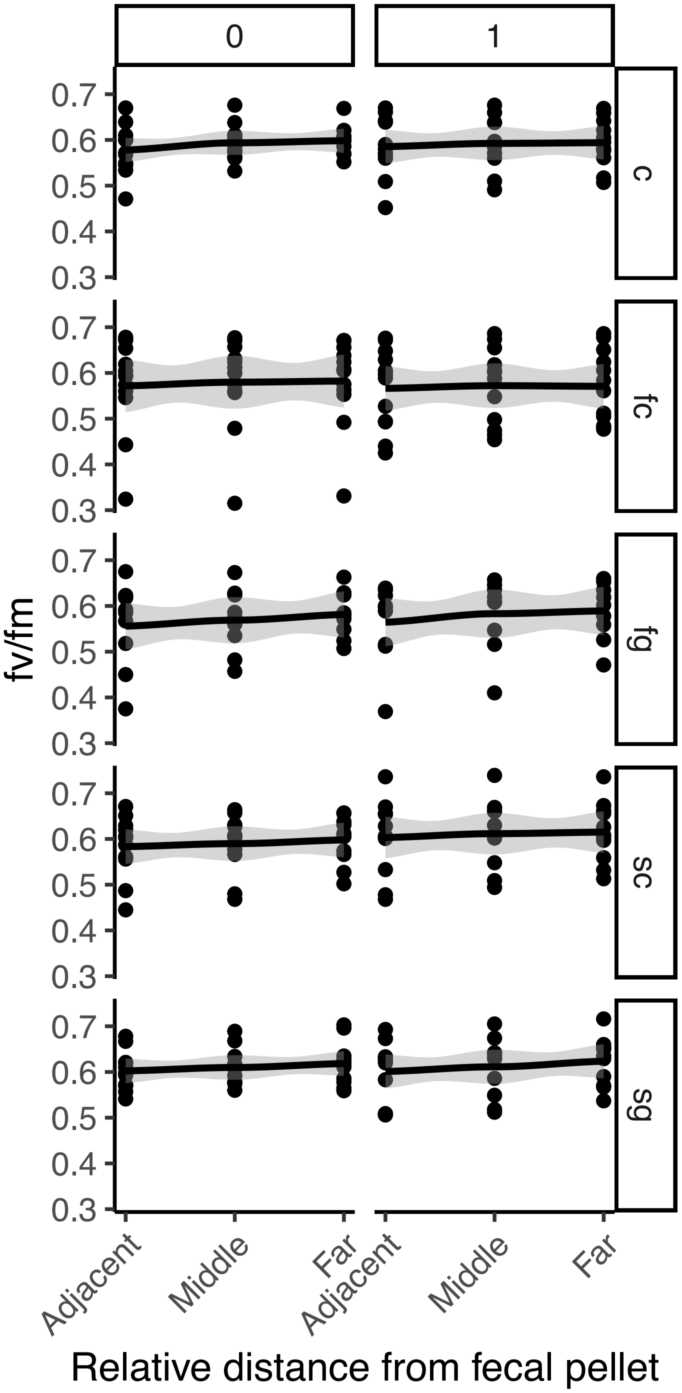


**Figure S2** Photosynthetic efficiency of coral fragments before (timepoint 0) and after (timepoint 1) fecal treatments were applied and removed. There was no effect of fecal treatment on coral photosynthetic efficiency (LM results: treatment*distance: Df=8, F=0.045, p=0.999; treatment: Df=4, F=2.27, p=0.065; distance: Df=2; F=0.53, p=0.588). C: Control; fc: fresh corallivore feces; fg: fresh grazer feces; sc: sterile corallivore feces; sg: sterile grazer feces.
